# Modelling co-development between the somites and neural tube in human Trunk-like Structures (hTLS)

**DOI:** 10.1101/2024.12.16.628661

**Authors:** Komal Makwana, Louise Tilley, Probir Chakravarty, Jamie Thompson, Peter Baillie-Benson, Ignacio Rodriguez-Polo, Naomi Moris

## Abstract

Human stem cell-based embryo models have opened new avenues of research in development by providing experimentally amenable *in vitro* systems. One of the features of embryo models is their multilineage differentiation, which allows the co-development of, and interactions between, tissues. Here, we utilise a human Trunk-like Structure (hTLS) model to explore trunk development. We show that hTLS have morphologically organised somites and a neural tube that form through self-organised, endogenous signalling including anteroposterior FGF, Wnt and Retinoic Acid (RA) gradients that modulate the fate of neuromesodermal progenitors. Comparison to an existing dataset from organogenesis-stage human embryos shows that hTLS cells approximate Carnegie Stage 13-14 (28-35 days post-fertilisation). The absence of a notochord leads to a dorsal identity, but exogenous exposure to smoothened agonist (promoting Sonic Hedgehog signalling) progressively ventralises both somites and neural tube in a dose-dependent manner. Furthermore, we identify endogenous signalling from the neural tube to the somites, which leads to medially localised *ALDH1A2,* and subsequent RA signalling from the somites to the neural tube, which leads to spontaneous neural progenitor patterning and *PAX6* expression. Together, our data highlight the value of modularity in embryo models which we leverage to explore human trunk co-development.

**Summary:** Using a 3D, stem cell-based model of human embryonic trunk development, we examine the interactions across somitic and neural tissues to better understand the dynamics of human trunk co-development.

---

In the embryo, tissues and organs are specified and mature alongside neighbouring tissues, using local interactions that are important for correct development and downstream function. For example, neighbouring tissues may provide positional information to cells through spatially localised cues, or they may provide signalling feedback that balances proportions of cell fates and promotes maturation to specific cell types. Inter-tissue interactions are also likely to be important for ensuring synchronised developmental timing across tissues, so that the embryo keeps a consistent pace during organogenesis. We refer to this inter- dependency between tissues as ‘co-development’, and believe this phenomenon is likely to be critical for fully understanding how tissues and organs form with implications for *in vitro* protocols aimed at generating specific cell types of interest in the future.

There is therefore a need to investigate organogenesis at the multi-tissue level, rather than focusing on a single tissue at a time. This has been challenging using embryos because signalling pathway convergence and overlapping communication across many tissues can make it difficult to identify specific reciprocal interactions between any two tissues of interest. Classical experimental embryology provides a route to explore tissue interactions in the development of functional organs through manipulations including explants, transplants and excisions (see for instance, ^1–8^). However, these approaches are not feasible in humans due to ethical and technical constraints, including limited long-term human embryo culture^9^.

Stem cell-based embryo models offer a practical alternative^10^. Their multilineage potential enables exploration of inter-tissue interactions, while bottom- up generation of specific tissues in the presence or absence of others can determine the necessity and sufficiency of these interactions^11^. Importantly, human pluripotent stem cell (PSC)-derived models provide scalability, experimental flexibility, and species-specific relevance that cannot be achieved with embryos alone. As such, these models hold promise for deciphering the regulatory logic of co-development during early organogenesis.

Several models of early embryogenesis have now been developed that allow researchers to examine different stages of human development. Human gastruloids are self-organised aggregates of pluripotent stem cells (PSCs) that spontaneously break symmetry, differentiate to all three germ layers, and undergo axial elongation and spatiotemporal gene expression along an anteroposterior axis^12^. While they are a valuable research tool to explore early cell fate decisions, including the differentiation to early paraxial mesoderm, they lack morphological complexity including somitic epithelialisation. Mouse gastruloids can be induced to form axially- organised morphological somites through embedding in an extracellular matrix gel^13^ and even form integrated somite- and neural tube-containing structures called Trunk- like Structures (TLS^14^). Likewise, three-dimensional human stem cell models of somitogenesis^15–17^ and caudal neural tube development^18,19^ have been described, as have structures that contain both neural and somitic mesoderm^20–22^, neuromuscular junctions^23,24^ and notochord progenitors^25^. These models showcase the remarkable modularity of stem cell-based embryo models to include or exclude particular tissues, in a building block-like manner. We reasoned that this could be harnessed to explore co-development, focusing particularly on the tissue interactions during axial extension in early embryogenesis.

The somites and the neural tube are the sources of trunk musculature, dermis, bone and spinal cord, respectively. Their development needs to be coordinated in order to form functional organ systems, for instance, specific neurons emanating from the spinal cord must innervate specific muscles in order to form circuits. Additionally, their close proximity next to one another during development makes it likely that general communication and feedback between the tissues enables co-development of both tissues. There is evidence that signals from the neural tube promote myogenesis ^3,5–7,26^ and prevent medial *Sim1* expression in the somites^2^, while somitic signals to the neural tube promote neurogenesis^8,27^.However, the exploration of these interactions in humans has been hampered for lack of appropriate experimental systems. Here we modify a somite-only axioloid protocol to develop robust and reproducible human Trunk-like Structures (hTLS) that could be directly compared with axioloids to explore trunk development in the presence or absence of the neural tube.

We find that, due to the absence of a notochord, the hTLS exhibit dorsal bias but exogenous activation of Sonic Hedgehog (Shh) signalling induces progressive ventralisation in a dose-dependent manner. Notably, even without a localized Shh source, we observed neural tube patterning orthogonal to the anterior-posterior (AP) axis, mediated by somite-derived Retinoic Acid (RA) signalling. This RA signalling was required for localized *PAX6* expression in the neural tube but not for initiating neurogenesis. Moreover, we found that the neural tube was able to induce *ALDH1A2* expression specifically in medial somites, a novel finding that we validated in mouse embryos, indicating that reciprocal interactions may promote somitic mediolateral identity. Additionally, in the presence of a developing neural tube, the development of the somites occurs at a pace that is comparable to that of the embryo. Our findings highlight the potential of stem cell-based embryo models to uncover the regulatory logic of co-development, and show that the modular nature of embryo models can offer insights into the spatial and temporal coordination of tissue interactions during early human organogenesis.

## Results

### Establishing an hTLS protocol

While mouse gastruloids embedded in low percentage Matrigel can develop organised epithelial somite structures^13,14^, human gastruloids had to be exposed to the Nodal signalling inhibitor, SB431542 (herein, SB43) during pre-treatment^12^, to generate segments when embedded in 20% Matrigel (Supplementary Figure 1a). These segments formed sequentially from the posterior, but they did not form rounded, epithelial structures characteristic of somitic morphology in the embryo. Instead, a protocol combining the Wnt/β-catenin activator CHIR99021 (herein, Chi) and SB43 at aggregation with embedding in 10% Matrigel enabled morphologically organised somitic tissue formation as axioloids^15^ (similar to ^16,17^). We adapted this protocol from induced pluripotent stem cells (iPSC) to human embryonic stem cells (hESCs; Supplementary Figure 1b) by adjusting the Chi concentration according to cell line requirements (see Methods). In doing so, we noticed that higher concentrations of Chi biased the cultures towards the mesodermal fate, as expected^28^, while lower concentrations enabled the co-formation of a morphologically organised neural tube alongside the somites (Figure 1a). These structures increased in somite number, elongated over 120h and displayed an organised somitic and neural tube morphology in 79% of aggregates (Figure 1b; Supplementary Figure 1c- h). Marker gene expression confirmed the presence of somitic (MEOX1+) and neural (SOX2+) identities with a pool of progenitors (SOX2+,TBXT+) at the posterior, and a presomitic mesoderm (PSM) domain (*TBX6+,MESP2+*) in the tailbud region (Figure 1c-d). As such, the structures were clearly similar to recently-described human trunk-like structures (hTLS)^20–22^, with the important distinction that the protocol could be ‘fine-tuned’ by adjusting the level of Chi to give somite-only or combined somite- and neural tube-containing structures, thereby allowing direct comparison between structures with the presence or absence of the neural tube.

**Figure 1:**
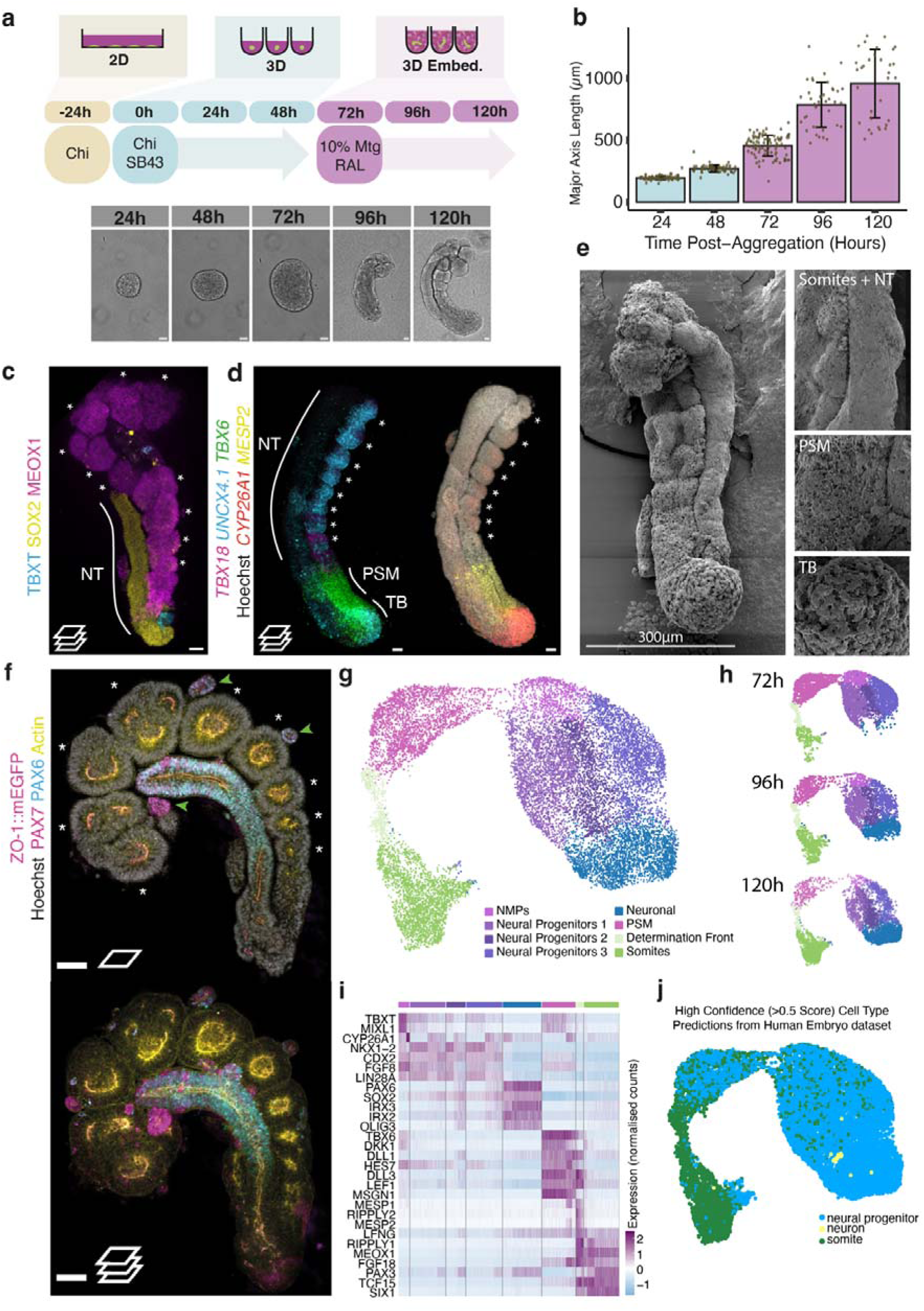
Establishing a human trunk-like structure (hTLS) model. **(a)** Schematic of hTLS protocol with representative brightfield images at each 24h timepoint. **(b)** Elongation of ZO-1::eGFP hTLS major axis length (mean±standard deviation(sd)). **(c)** Projection of immunofluorescence imaging of hTLS at 120h. NT, neural tube; * indicates somites. Scale bar, 50 μm. **(d)** Projected images of whole mount, *in situ* hybridization chain reaction (HCR) staining of hTLS at 120h. **(e)** Scanning electron micrograph of a hTLS at 120h. Scale bar, 300 μm. **(f)** Confocal section (top) and projection (bottom) of immunofluorescence imaging of a ZO- 1::eGFP hTLS at 120h. Green arrows highlight clusters of PAX7+ cells. Scale bars, 100 μm**. (g)** Single cell transcriptomic UMAP of hTLS development. **(h)** Temporal spread of samples collected from 72h, 96h and 120h timepoints. **(i)** Marker gene expression for each identified cluster. **(j)** Projection and subsequent label transfer of high confidence cell identities from a published human embryo dataset^37^. Chi, Chiron (CHIR99021); Mat, Matrigel; SB43, SB431542; RAL, retinal; TB, tailbud; PSM, presomitic mesoderm; NMP, Neuromesodermal progenitor. Scale bars, 50 μm, unless otherwise stated.

Like several other *in vitro* models of somitogenesis^15,21^, we noted that the morphological organisation of hTLS was critically dependent on embedding in Matrigel and exposure to retinal, since time-matched structures without these had some degree of gene expression organisation along an AP axis, but no or little morphological organisation of somites or neural tube (Supplementary Figure 1i-j). We were also able to adapt the protocol to a total of 7 cell lines (from 3 parental backgrounds), including both hESC and iPSCs, which showcases the versatility of the protocol beyond intrinsic line-to-line variability (Supplementary Figure 2; Methods).

### Characterising hTLS formation

To better understand the morphological organisation of hTLS, we performed Scanning Electron Microscopy (SEM) to visualise the surface morphology of hTLS. This showed clear cell compaction differences, with loosely arranged cells in the posterior tailbud, medium compaction in the PSM, and tightly compacted epithelialisation in both the neural tube and somite domains (Figure 1e), in keeping with descriptions in vertebrate embryos^29,30^. Likewise, hTLS generated with a ZO1::mEGFP line showed that both the somites and neural tube had internal ZO1- and Actin-rich lumina, with cells organised as epithelia in both tissues (Figure 1f). We therefore wanted to better understand how these epithelial tissues formed in hTLS.

This question is particularly interesting for the neural tube since it is known to form through two modes of morphogenesis in mammals: primary and secondary. Primary neurulation involves the folding of the neural ectoderm^31^ and has been extensively studied because malformations are the leading cause of neural tube defects (NTDs) in humans^32^. By contrast, secondary neurulation - when neural cells condense to form rods that later epithelialize to form a contiguous tube - is much less understood since it accounts for only the caudal-most portion of the axis^33^. Exactly how secondary neurulation occurs in humans, how the two modes are coordinated, and whether the human embryonic neural tube is branched at early stages or contiguous^34^ are still open questions in the field. Using the ZO1::mEGFP cell line, we sometimes saw multiple or branched neural tubes, particularly at the posterior pole (Supplementary Figure 1k). With live imaging we observed epithelial cysts that progressively merged to generate a contiguous tract (Supplementary Figure 1l & Supplementary Movie 1). Therefore, it seems that the mode of neural tube formation in hTLS structures is more similar to secondary neurulation.

To further characterise the hTLS over time, we collected samples at 72h, 96h and 120h and performed 10x single cell RNA sequencing. We could detect populations including a neuromesodermal progenitor (NMP) cell state (expressing *SOX2* and *TBXT*), PSM (*TBX6, HES7, DLL1*), determination front (*RIPPLY1/2, MESP1/2*), somitic (*MEOX1, TCF15*), and neural (*SOX2, PAX6*) clusters (Figure 1g, Supplementary Figure 3a-e). Cells from the early timepoints mostly populated the progenitor populations while those from later timepoints populated the more mature states (Figure 1h). In addition, we detected sub-clusters including a small population of endothelial cells (*KDR+, ETV2+;* Supplementary Figure 3f) that match previous reports of a somite-derived endothelial population in mouse TLS^14^, human axioloids^15^ and embryos^35,36^. Additionally, we observed a small population of more mature neural cells (*ONECUT2, ELAVL3*; Supplementary Figure 3g) indicating that neurogenesis is beginning by 120h in hTLS. By contrast, we saw no evidence of cell fates correlating to notochord, non-neural ectoderm, neural crest, endoderm, lateral plate mesoderm or others, corroborating the observation that the hTLS contain only neural tube and somitic progenitors and downstream lineages (Figure 1i). Bioinformatic projection onto an existing axioloid dataset^15^ confirmed that the hTLS contain neural cell types that are not present in the somite-only version (Supplementary Figure 4) indicating that, while comparable, hTLS are indeed distinct entities from axioloids.

To further examine whether our annotated clusters had equivalents to their corresponding *in vivo* states, we projected our dataset onto a human embryonic atlas of stages ranging from Carnegie Stage (CS) 12 to CS16 that incorporates all major cell types at early organogenesis stages^37^. Our data clearly mapped to the somitic and neural progenitor populations of this dataset, once again highlighting that the *in vitro* populations observed in hTLS match the expected cell types in the human embryo (Figure 1j; Supplementary Figure 5a-b). Interestingly, when we performed label transfer for our dataset, the vast majority of cells were called as CS13-14 equivalent (day 28-35 of human development; Supplementary Figure 5c-g) which is significantly later than that predicted for human gastruloids (approximately CS8-9; day 17-21 of human development)^12^, indicating the significant step forward achieved by the hTLS protocol.

### Axial elongation in hTLS is maintained by an NMP population

We next examined the spatial and temporal regulation of Hox gene expression, as these genes are known to be critical in the regulation of axial identity along the anteroposterior (AP) axis of vertebrate embryos^38^. We observed expression of *HOXC6/8* along the full AP axis of hTLS, while *HOXC10* was only present in the posterior portion (Supplementary Figure 6a). Temporally, there was progressive expression of HOX gene paralogues across the timepoints sampled (Supplementary Figure 6b) consistent with spatiotemporal regulation. Together with the lack of expression of HOX11-13, this suggests that the hTLS model approximates the thoracic and lumbar regions of the AP axis, but not the sacral or caudal-most regions^39^. However, given that the termination of axial elongation is thought to be triggered by expression of HOX13 in embryos^40,41^ which is not present in hTLS, we wondered what was mediating the extension and then termination of axial elongation in our hTLS model.

In the embryo, axis extension is supported by a population of neuromesodermal progenitors (NMPs): bipotential progenitors that give rise to both the neural and somitic cell fates (Figure 2a)^42–44^. One basic characterisation of NMPs is co-expression of *TBXT* and *SOX2*^45^ which is thought to underly their bipotency, and we observed cells co-expressing these markers in the posterior of hTLS (Figure 2b). Using the RUES2-GLR cell line^46^, which contains TBXT-mCerulean and SOX2- mCitrine fluorescent reporters, we live-imaged hTLS under conditions that promoted co-development of neural tube and somites (hTLS), or conditions that biased towards somite-only (Chi high conditions) or neural tube-only fates (Chi low conditions; Figure 2c). We observed an early bias (between 78-90h) towards SOX2- only or TBXT-only expression in neural tube-only and somite-only structures respectively, with co-developing structures maintaining a TBXT and SOX2 co-expressing domain for the longest duration, consistent with the role of NMPs in contributing to both fates.

**Figure 2:**
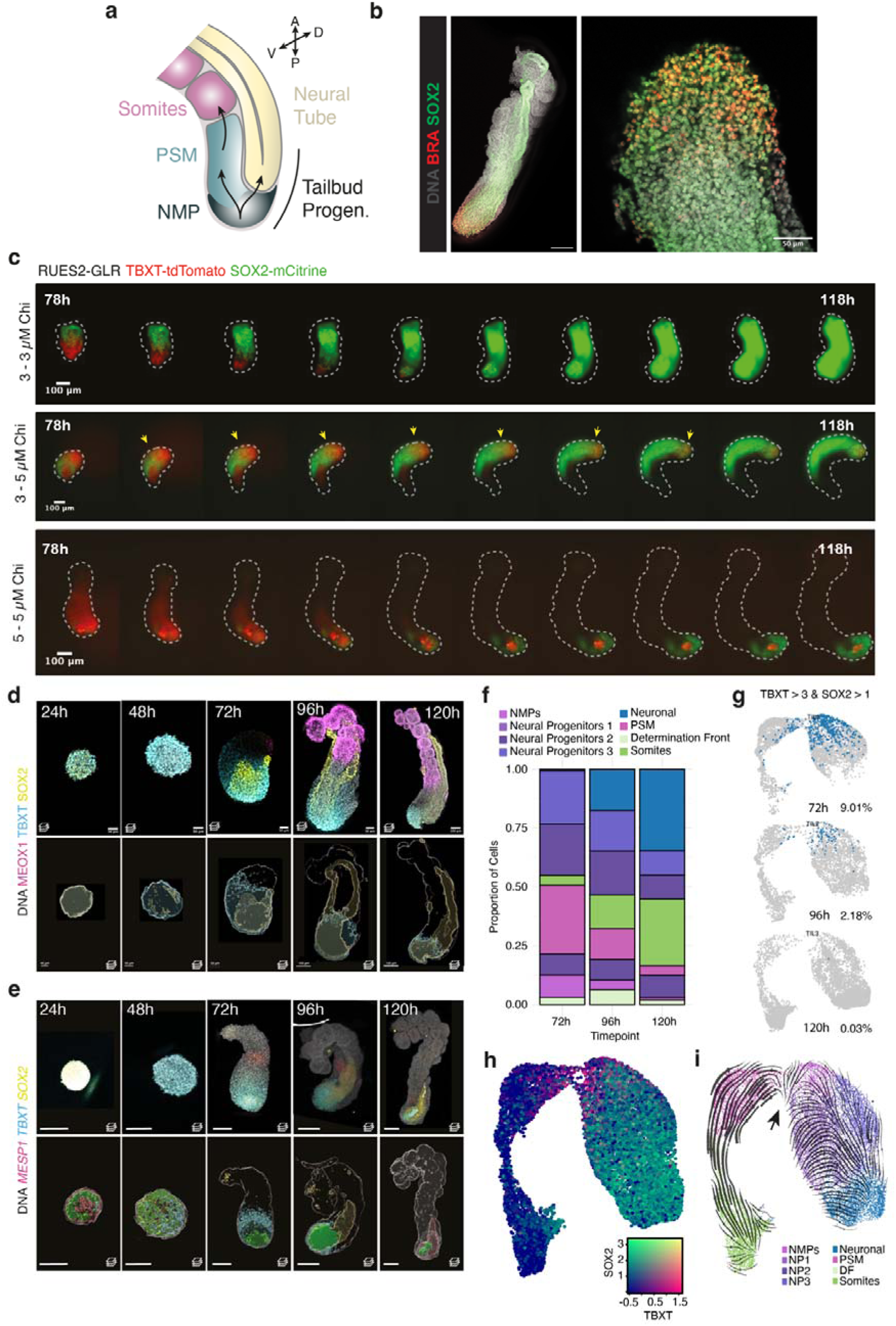
Dynamic neuromesodermal progenitor (NMP) population in hTLS. **(a)** Schematic of NMPs, that contribute to both somites and neural tube tissues. **(b)** Co-staining of TBXT and SOX2 expression. **(c)** Representative images of RUES2-GLR hTLSs imaged at 4h intervals over a period of 40 hours. Dotted line outlines the brightfield images to show full hTLS structure. **(d-e)** Staining of hTLS (top) and Imaris-rendered regions of positive expression of TBXT (blue), SOX2 (yellow), and dual TBXT/SOX2 co-expression (green), by immunofluorescence **(d)** and HCR **(e)**. Z-step, 2 µm. Scale bars, 100 µm. **(f)** Proportion of cells within each cluster identity over time. **(g)** Percentage and location of double positive (TBXT>3 and SOX2>1 normalised counts) cells over time, 72h (top), 96h (middle) and 120h (bottom). **(h)** Co-expression of SOX2 and TBXT for hTLS cells projected onto a UMAP. **(i)** RNA velocity map, showing vectorial directions of predicted differentiation. Black arrow indicates a branchpoint within the annotated Neuromesodermal progenitor (NMP) cluster. NMP: Neuromesodermal progenitors; NP: Neural Progenitors; PSM: Presomitic mesoderm; DF: Determination front; D, dorsal; V, ventral.

However, the NMP domain showed a clear decrease in volume over the hTLS time-course, as was confirmed by fixing and staining at the mRNA and protein level (Figure 2d-e) and by our single cell RNA-sequencing data (Figure 2f-h). Indeed, RNA velocity analysis showed evidence of bidirectional fates predicted from this population (Figure 2i), and trajectory analysis rooted in the NMP cluster predicted endpoints in both somites and neural tube populations (Supplementary Figure 7). Together this data suggests that hTLS contain an NMP population capable of bipotential cell fate differentiation, and its gradual depletion may be responsible for the resultant maximum axial length and number of somites observed in hTLS *in vitro*.

### Endogenous signalling patterns in hTLS

In vertebrate embryos, the sequential addition of somites from the posterior unsegmented region is determined by a combination of local intrinsic oscillations of gene expression and a global signalling landscape that leads to coordinated and organised somite formation^47^. Using a HES7::Achilles iPSC line^48^, we observed oscillatory expression in the PSM of hTLS, that moved posterior to anterior and stopped at the site of emergent somite formation (Figure 3a-b; Supplementary Movie 2), as has been described in embryos^49,50^. The time-period of these oscillations approximated an average of 373 mins (6.2 hours; Figure 3c) which is longer than *in vitro* measurements made in alternative human systems (∼5 hours in Somitoids^16^ and Axioloids^15^, 4.6 hours in Segmentoids^17^) but closer to the predicted period from human embryos (7.2 hours)^51^, raising the possibility that the presence of the neural tube might slow an intrinsic somitic clock to coordinate the developmental of the two structures. Further research would be necessary to investigate the mechanism of this influence.

**Figure 3:**
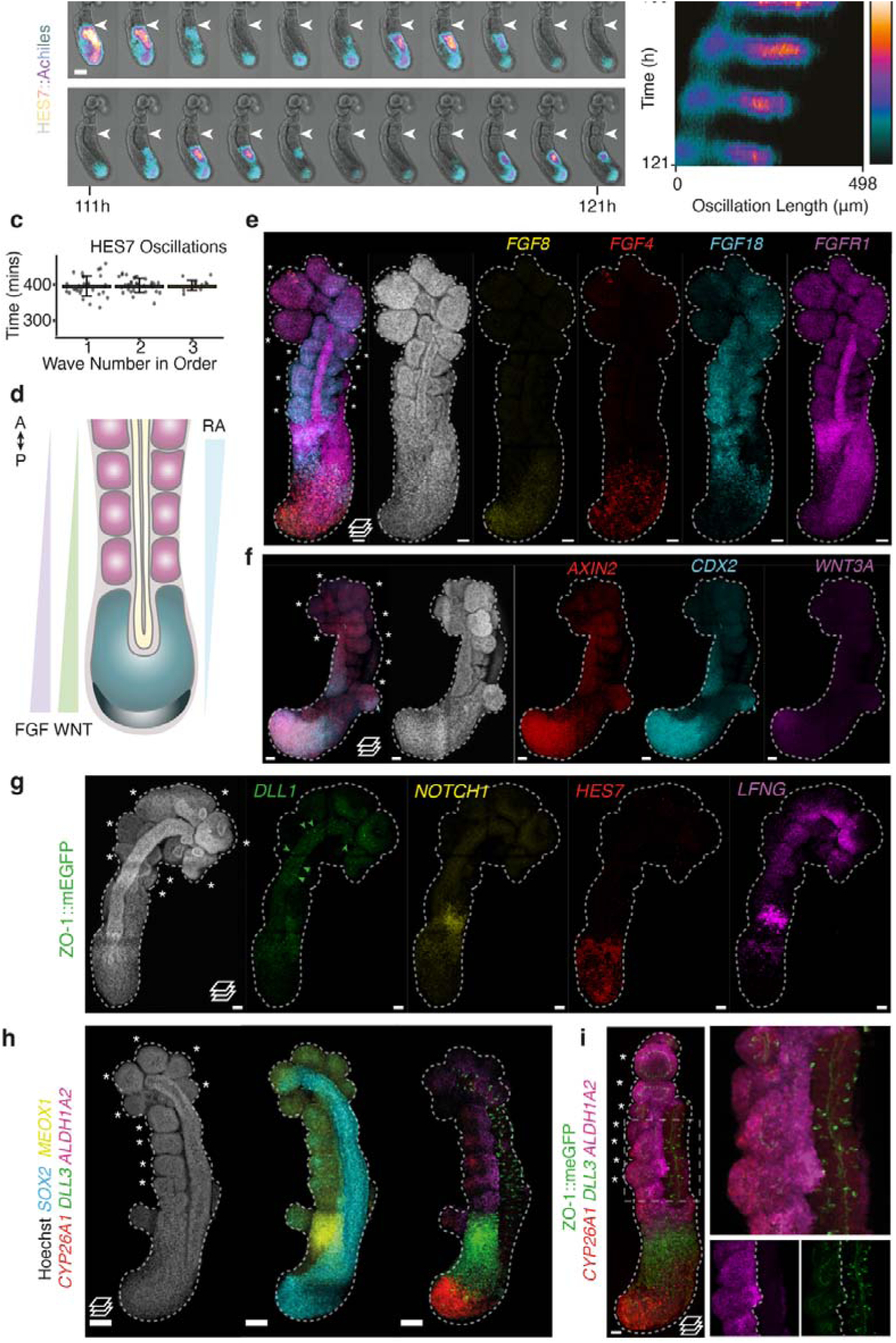
Characterising signalling in hTLS models. **(a)** Representative stills from live imaging of HES7-Achilles hTLS at 20-minute intervals. White arrows denote the same boundary, showing posterior shift of the wavefront. Scale bar, 100 µm. **(b)** Kymograph of Achilles oscillations from HES7::Achilles hTLS represented in (a). **(c)** Time intervals of HES7 oscillations from peak-peak across 21h period (mean± standard deviation(sd)). **(d)** Schematic of anterior-posterior signalling gradients across the embryo axial length. **(e-i)** Projected images of HCR staining of hTLSs at 120h for components of the **(e)** FGF, **(f)** WNT, **(g)** Notch and **(h)** RA signalling pathways (Scale bars, 100 μm). (**i)** Localised mediolateral signal of *ALDH1A2* in somites and punctate signal of *DLL3* through neural tube. Dashed box shows location of enlarged region. Scale bars, 50 μm unless otherwise stated; Green arrows, *DLL1* signal in the neural tube; * indicates somites; Dashed line outlines hTLS; A, anterior; P, posterior; RA; retinoic acid.

We then turned our attention to the signalling landscape of the hTLS along the AP axis. In the embryo, the signalling gradients primarily responsible for the wavefront are thought to be FGF and Wnt signalling at the posterior, and Retinoic Acid (RA) more anteriorly at the level of the somites (Figure 3d). We observed *FGF8* and *FGF4* in the posterior region^52,53^; *FGF18* expression in the first 3-5 newly-formed somites, the determination front and, to a lesser extent, the anterior PSM^54^; and *FGFR1* in the neural tube and the determination front (Figure 3e)^52,55^. We also saw posterior expression of *WNT3A* and *AXIN2* in the tailbud (Figure 3f), consistent with a posteriorly localised Wnt signalling gradient^56–59^.

Likewise, we observed expression of Notch signalling ligands *DLL3* and *DLL1* strongly in the anterior PSM and determination front and in a salt-and-pepper organisation in the neural tube (Figure 3g-h), matching descriptions in the embryo, where a Notch-dependent mutual inhibition mechanism leads to neural differentiation^60–62^. In addition, expression of *LFNG* and *NOTCH1* in the anterior PSM and newly formed somite are consistent with the role of Notch signalling in mediating the dynamic control of somitogenesis^63^ (Figure 3g).

The crucial anterior signalling gradient that opposes the Wnt and FGF gradients is Retinoic Acid (RA), which inhibits posterior FGF signalling and fine-tunes the progenitor-to-differentiated neuron gradient along the neural tube^8^. Typically, ALDH1A2 (an alcohol dehydrogenase involved in the synthesis of RA) and CYP26A1 (which degrades RA) have been used together as a proxy for RA signalling status in chick^64^ and mouse embryos^65^. In hTLS, *CYP26A1* is strongly expressed in the most posterior tip of the tailbud, with *ALDH1A2* expressed in the somites (Figure 3h-i) indicative of a ‘source and sink’ pattern creating an AP gradient of RA signalling. As such, it is likely that self-organised signalling along the AP axis provides endogenous signals that establish the clock-and-wavefront mechanism of somitogenesis in hTLS.

### Exogenous Shh defines the dorsoventral identity of hTLS structures

In the embryo, signals from the surrounding tissues including the notochord, lateral plate mesoderm and surface ectoderm, are all integrated to provide positional information to cells within the neural tube and somites^66^. In embryo models, the presence or absence of such tissues can allow us to investigate the extent of positional information in a simplified system. For instance, Shh signals from the notochord play a key role in setting up the dorsoventral (DV) axis in both the neural tube and somites (see Sagner and Briscoe ^67^ for recent review). We therefore wondered what DV identity our hTLS structures assumed in the absence of these tissues.

Based on transcriptomic data analysis, we found that the hTLS neural tube has a dorsal identity (*PAX6, OLIG3* and *IRX3)*, with no detectable ventral (*OLIG2, NKX2-2/8*), floor plate (*FOXA2, SHH*) or roof plate markers (Figure 4a-b). Indeed, comparison of the hTLS neural clusters with human embryonic spinal cord data^68^ showed that they aligned most strongly to progenitor dorsal cell identities, excluding the roof plate (Supplementary Figure 8a).

**Figure 4:**
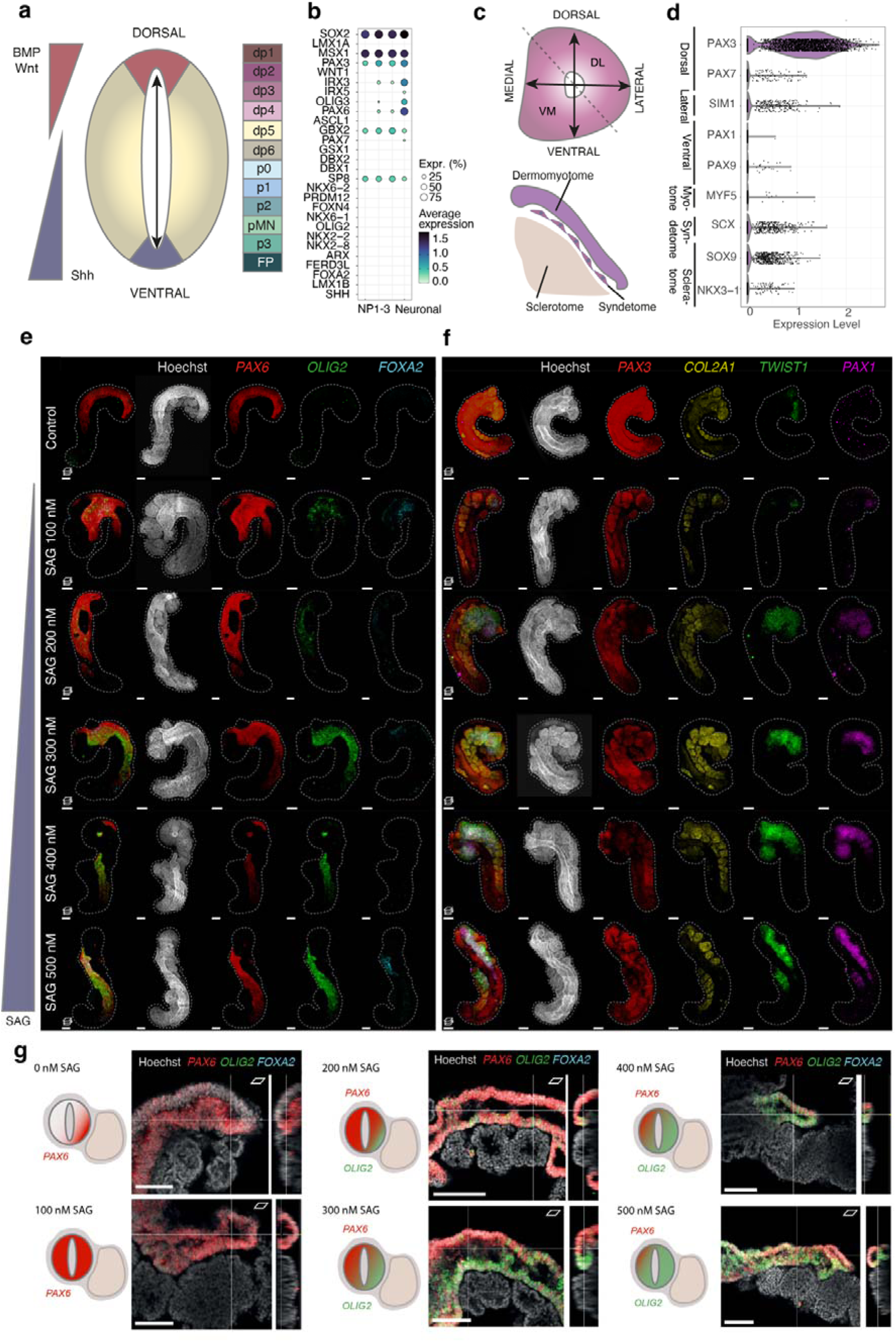
Dorsoventral patterning in hTLS structures. **(a)** Schematic diagram of the signalling gradients (left) and subsequent dorsoventral (DV) patterning of the neural tube (middle), leading to discrete neural populations (right). **(b)** Gene expression of neural populations in hTLS, ordered from top to bottom by expression along the DV axis in embryos. **(c)** Schematic diagram of the DV and mediolateral axes of the developing somite (top) and subsequent tissue maturation (bottom). **(d)** Gene expression signature of somites in hTLS. **(e-f)** Projected images of 144h hTLS treated with increasing concentrations of SAG (sonic hedgehog pathway activator) and HCR stained for **(e)** neural and **(f)** somitic DV markers. Dashed line outlines hTLS boundaries. Scale bars, 100 µm. **(g)** Schematic diagrams (left), XY images (middle) and YX optical slices (right) showing HCR gene expression in the 144h hTLS neural tube with increasing SAG concentrations. Scale bars, 150 µm for SAG 200 nM condition and 100 µm for all others.

Likewise, the somitic identity (Figure 4c) of hTLS show expression of pan- somite markers (*TCF15*, *MEOX1, FOXC2)*, dorsal markers of dermomyotome (*PAX3, PAX7, SIX1, SIX4, EYA1*) and syndetome (*SCX*; Supplementary Figure 3c,h), with very few late dermatome or myotome markers (*MYF5, MYOD, MYOG;* Figure 4d). We observed few cells expressing markers of ventral somitic identity, the sclerotome, with only a handful expressing detectable *PAX1* or *PAX9* (Figure 4d). Together, this suggests that the lack of notochord as a source of Shh signalling leads to an overall dorsalisation of both the somites and neural tube in hTLS.

To test whether hTLS do still have the capacity to ventralise, we exposed the structures to varying concentrations of Smoothened Agonist (SAG) which promotes Shh signalling, as well as to the small molecule Cyclopamine that inhibits Shh signalling^69^ as a control (Supplementary Figure 8b). Upon SAG addition we observed increasing ventralisation of hTLS structures in a dose-dependent manner, with increased expression of ventral neural tube (*OLIG2*, *FOXA2*) and ventral somite (*PAX1, TWIST1*) markers (Figure 4e-f). Additionally, the morphology of hTLS appeared to be altered under increasing SAG exposure with less compacted anterior somites (Supplementary Figure 8c), suggesting that epithelial-to-mesenchymal transition (EMT) associated with ventral (sclerotome) fates could be occurring, while cyclopamine had no observable effect on hTLS (Supplementary Figure 8d). However, the dose-dependent ventralisation of hTLS exposed to Shh signalling was not simply all-or-nothing. Surprisingly, we observed that at high concentrations of SAG exposure, not only did populations of *OLIG2+(PAX6-)* and *PAX6+(OLIG2-)* appear in the neural tube, but these were organised across an axis orthogonal to the AP axis (Figure 4g). In the embryo, it is known that negative feedback between transcription factors acts to establish discrete domains of neural progenitor identity along the continuous Shh gradient^70^, and the same is likely happening in the hTLS model. However, since the hTLS are exposed to a single, exogenous concentration of SAG throughout (rather than a localised source) the appearance of a consistent axis was surprising. In this case, we reasoned that the juxtaposition of the somites next to the neural tube may establish some aspects of an axis in the neural tube, since the *OLIG2+(PAX6-)* domain was consistently localised adjacent to the somites (Figure 4g). We therefore set out to investigate the nature of this communication between somites and neural tube.

### Signalling interactions between tissues

One of the primary benefits of a co-developing system like the hTLS is the ability to use it to probe the interactions between tissues, which would otherwise present challenges in an embryonic context. Beyond the difficulties of accessing human embryonic material at these stages of development, the complexity of the signalling landscape – with signals emanating from diverse tissues and organs, across multiple spatial axes, and dynamically changing over time – makes it complicated to unravel the contribution of a single tissue. In this case, we were interested in asking what the signalling interactions are between the neural tube and somites, in the absence of all other tissues (Figure 5a).

**Figure 5:**
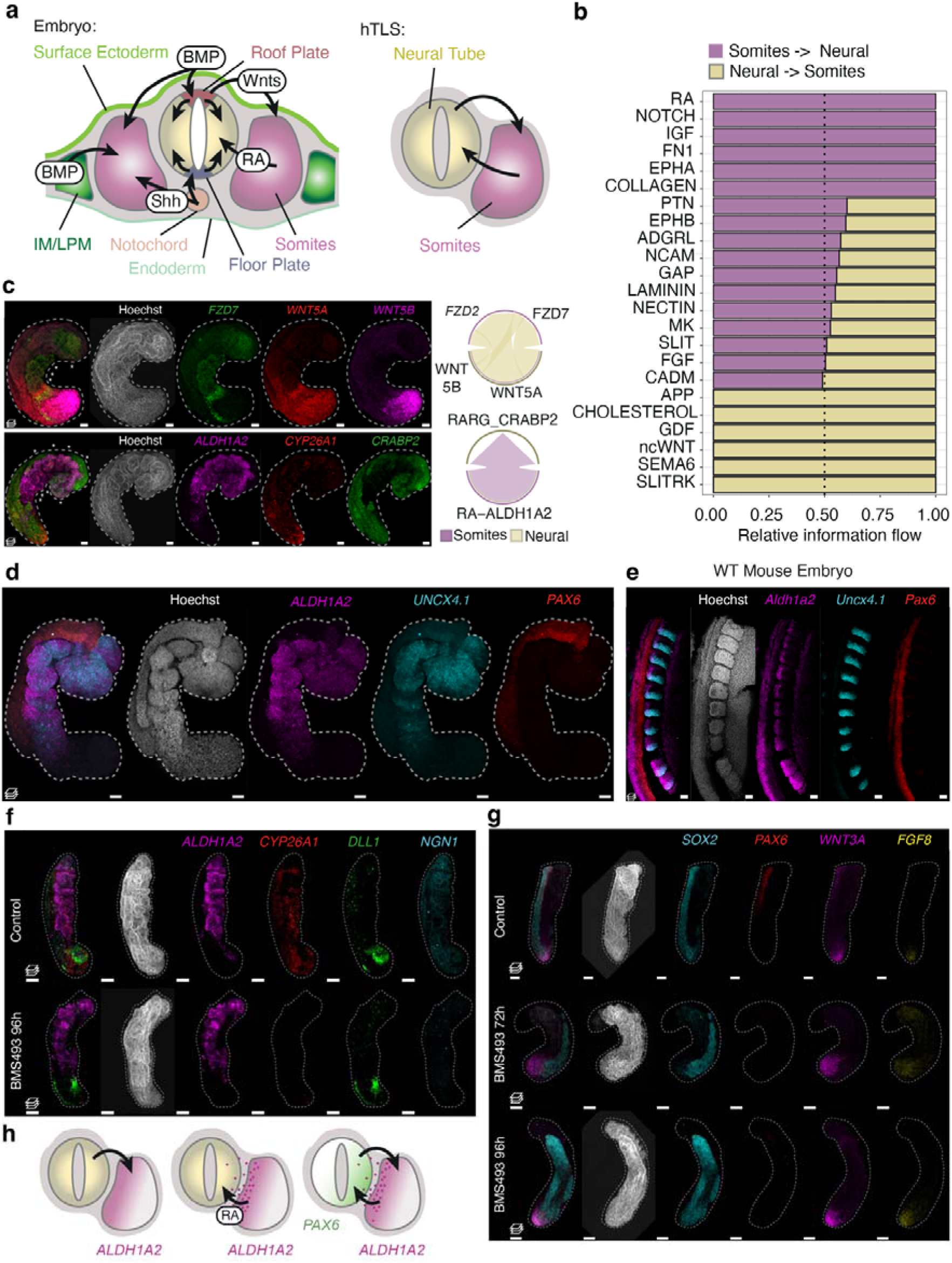
Interactions between somites and neural tube in hTLS and embryos. **(a)** Schematic diagram of the reciprocal signalling environment of the embryo (left) and hTLS (right). **(b)** Relative information flow from CellChat^71^ analysis, identifying unique signalling interactions between the somite cell cluster and neural clusters (combined Neural progenitor populations 1-3 and Neuronal clusters). Dotted line indicates equal information flow = 0.5. **(c)** Localised expression of non-canonical Wnt (top) and RA (bottom) signalling components by HCR in 120h hTLS (left), as predicted from CellChat analysis (right). **(d-e)** Somite-specific *ALDH1A2* expression adjacent to the neural tube, in 120h hTLS **(d)** and E9.5 mouse embryo **(e)**. **(f-g)** Projected images of hTLS at 120h following retinoic acid (RA) inhibition at 72h or 96h (BMS493, RA inhibitor). Structures were examined for differences in RA signalling (*ALDH1A2, CYP26A1)* and neurogenesis (*DLL1*, *NGN1*; **f**) or neural (*SOX2*, *PAX6)* and posterior tailbud (*WNT3A*, *FGF8)* gene expression **(g)**. **(h)** Schematic diagram of proposed reciprocal signalling occurring between somites and the neural tube of hTLSs. Signals from the neural tube bias *ALDH1A2* expression in the somites towards the neural tube (left), followed by subsequent RA signalling from somites to neural tube, leading to PAX6 expression biased towards the somitic side (right). RA, retinoic acid; IM, Intermediate mesoderm; LPM, Lateral plate mesoderm. Scale bars, 50 μm; * indicates somites; Dashed line outlines hTLS.

We turned first to unbiased, computational approaches to interrogate the putative signalling ligand-receptor interactions predicted between the two tissues. Building on the expression profile of ligands and receptors of known pathways (Supplementary Figure 8e), we used CellChat^71^ to identify statistically significant, unidirectional interactions specifically from the somitic to the neural tissues (combining neural progenitor populations 1-3 and neuronal clusters), or the inverse interaction. This identified several putative signalling interactions, including RA, Notch, IGF, Fibronectin and EPHA signalling from somites to neural tissue, and Cholesterol, GDF, non-canonical Wnt, SEMA6 and SLITRK pathway signalling from neural to somitic tissues (Figure 5b; Supplementary Figure 8f-g).

We validated the expression signatures of somite-derived RA and neural tube-derived non-canonical Wnt signalling by examining the localisation of the gene corresponding to an RA-binding protein, *CRABP2* (alongside *ALDH1A2* and *CYP26A1),* which was expressed in the neural tube as predicted (Figure 5c), and by expression of ligands *WNT5A* and *WNT5B* in the neural tube and receptor *FZD7* in the somites (Figure 5c). We therefore hypothesised that signalling between the two tissues in hTLS is responsible for setting positional identity and organising axis formation.

### Mediolateral ALDH1A2 expression in somites

An additional surprise came on closer examination of the somites, where we noticed that *ALDH1A2* was not uniformly localised across the somite, but instead was most strongly expressed adjacent to the neural tube (Figure 5d). This is in direct contrast with somite-only axioloid structures, where ALDH1A2 was shown to instead have an AP polarity^15^, consistent with our own experiments in axioloids (Supplementary Figure 9a). This hinted that the presence of the neural tube may bias the localisation of *ALDH1A2* expression to the neural tube-adjacent side, with important implications for the mechanism of medio-lateral polarisation in somites. Since we are not aware of this phenomenon being previously reported, we examined the same gene expression in E9.5 mouse embryos (Figure 5e) and likewise observed a clear medial bias of *ALDH1A2* expression in the somites (Supplementary Movie 3). This suggests that the neural tube biases the localisation of ALDH1A2 to the medial somite domain in both embryos and models, which may in turn localise the RA signalling exposure towards the neural tube.

### Somite-to-neural tube RA signalling in hTLS

We therefore turned our attention to the downstream role of localised *ALDH1A2*. In the embryo, RA signalling from the somites to the neural tube is thought to have three main functions: (i) AP axis organisation and morphogenesis of the neural tube, (ii) promoting cellular neurogenesis, and (iii) mediating the dorsoventral patterning of neural tube populations (see ^72^ for review). At subsequent stages of development, additional roles include coordination of neural crest cell specification^1^, interneuron specification^73^, establishment of the lateral motor column neurons^65,74,75^ and maintaining the bilaterality of the somites^76^. It has also been shown that the FGF pathway antagonises RA signalling, and that additional interplay with Wnt and Shh signalling might further refine the role of RA^8,77–79^. Indeed, joint exposure of neural progenitors to RA and FGF is sufficient, even in the absence of Shh signalling, to induce motor neuron differentiation^79^. In addition, in the absence of RA signalling, there is an expanded domain of ventral gene expression in the neural tube^80^, suggesting it has a dorsalising influence on the neural tube.

To test whether similar effects were also occurring in hTLS structures, we prevented RA signalling either by removal of the exogeneous retinal or by exposure to the pan-RA receptor (RAR) inhibitor, BMS493^81^. Following early (72-120h) exposure to BMS493, hTLS structures were morphologically impaired, with a smaller overall area, and were composed of fewer organised somites, smaller neural tube and increased tailbud size (Supplementary Figure 9b-c). This is consistent with the role of RA signalling in axial organisation and morphology, as *Raldh2^-/-^* mouse embryos die mid-gestation with aberrant morphology that precludes turning and includes a truncated posterior axis, small somites and an irregularly folded neural tube^82^. However, when hTLSs were exposed to BMS493 at later timepoints (96- 120h) there was no significant effect on morphometrics measured (Supplementary Figure 9b-c) suggesting that at this point, morphology had already been established and blocking the RA signalling might specifically affect the ongoing interactions between somites and neural tube. Indeed, hTLS structures exposed to BMS493 at either timepoint showed much lower or no *CYP26A1* in the neural tube, in contrast to control structures (Figure 5f; Supplementary Figure 9d) consistent with published data from chick embryos^80^. We therefore further investigated the proposed role of somite-derived RA signalling on the neural tube focussing on the latter timepoint, including its proposed role in promoting neurogenesis and setting up DV identity.

To investigate the role of somite-derived RA on early neurogenesis, we examined markers expression in hTLS structures, including *DLL1* and *NGN1* (Figure 3f; Figure 5f). Upon RA signalling inhibition, we could still observe *DLL1*+ cells in the neural tube, comparable to control levels (Figure 5f). This is interesting because in Vitamin A deficient (VAD) quails that lack the precursor to retinoic acid, retinal, *Delta- 1* expression is depleted from the caudal neural tube in 9-19 somite-stage embryos^8^, but in the *Raldh2*^-/-^ mouse model, *Delta1* expression is comparable to wildtype at the 14-15 somite stage^77^. This suggests that there might be differences between species in how RA signalling inhibition affects the neurogenesis cascade. As such, the observation that hTLS show RA-independent neurogenesis indicates that human development is likely to be more similar to the mouse than the avian embryo in this regard.

Finally, we examined the role of somite-derived RA signalling on patterning in the neural tube. It is known that *Raldh2^-/-^*mice and VAD quails have depleted expression of *Pax6* in the neural tube compared to wildtype embryos^83,80^. We therefore examined the expression of *PAX6* in the hTLS with late RA signalling inhibition and observed a substantial depletion of this neural marker from the neural tube structure (Figure 5g). Conversely, addition of all-trans Retinoic Acid (ATRA) onto hTLS structures increased the expression of *CYP26A1* in the neural tube, with no obvious difference in *DLL1* expression, and with broad expression of *PAX6* across the whole neural tube (Supplementary Figure 9d-e). Taken together, this shows that somite-derived RA signalling is required for *PAX6* expression in hTLS and, given the neural tube patterning observed (Figure 4), makes it likely that RA signalling is responsible for establishing this patterning. Similar Shh-independent RA signalling effects on neural progenitor identity have been shown in chick embryos^73^, but not in a human system.

Taken together, these data show that interactions between tissues include initial signalling from the neural tube to the somites, resulting in the medial somitic localisation of ALDH1A2 adjacent to the neural tube which, in turn, leads to localised RA signalling from somites to the neural tube that promotes *PAX6* expression specifically at the side adjacent to the somites (Figure 5h). Integration with the Shh pathway leads to progressive ventralisation while maintaining the self-organised patterning in the neural tube established by localised signalling from the somites. This reciprocal signalling between somites and neural tube may therefore play an important role in ensuring that correct axial patterning is coordinated between tissues and shows that elements of a mediolateral or dorsoventral axis can be established, even in the absence of well-known signalling centres such as the notochord or lateral plate mesoderm.

## Discussion

The hTLS model presented here showcases the development of the thoracic and lumbar trunk in a human system, which would not otherwise be possible given the inability to culture human embryos to this relatively late stage in culture or to study dynamic development in samples from terminations. As a result, the hTLS model serves as a platform to perform experiments that would otherwise be technically intractable, with the scale and experimental flexibility afforded by pluripotent stem cells. Complementary experiments using rare and highly valuable human embryonic material could validate predictions made from such *in vitro* models, thus yielding new insights into how the human embryo employs specific mechanisms for tissue co-development.

Our system has revealed a number of features associated with development of the paraxial mesoderm and neural tube. Interestingly, although hTLS represent the thoracic and lumbar region, the neural tube structures develop through a process similar to secondary neurulation, which is characteristic of the tail region^31^. This suggests that, in the embryo, the mode of neurulation is dependent on the early morphology at gastrulation, including the flat, epiblast-generated neuroectoderm, and since hTLS bypass these morphological stages they utilise secondary neurulation mechanisms instead. This may also help explain why different species evolutionarily rely on primary and secondary neurulation to different extents^84^, as a function of their early morphological constraints. It would be interesting to explore whether this substitution in the mode of neurulation plays a role on the cell biology or fate decisions in the neural tube in follow-on research.

However, the most significant findings from this work concerns signalling interactions between developing paraxial mesoderm and the neural tube. For instance, the hTLS model revealed medially localised *ALDH1A2* in the early somites of both hTLS and E9.5 mouse embryos, which to our knowledge had not previously been described. It could be that this early organised gene expression is necessary for subsequent differentiation into the epaxial and hypaxial muscles^85^, and may represent the earliest distinction of mediolateral identity in the developing somites. It also proves that neural tube-derived signals are *sufficient* to initiate mediolateral patterning in the somites, even in the absence of the intermediate or lateral plate mesoderm that would normally be localised laterally. The simplistic, modular nature of the hTLS model therefore provides enormous flexibility for experimental design, allowing us to systematically investigate the influence of particular tissues on the development of others.

We also identified spontaneous patterning of the neural tube orthogonal to the AP axis, based on somite-adjacent localization. We showed that this was due to RA-induced signalling emanating from the somites and leading to differential gene expression along the neural tube-to-somitic axis. While this is more likely to reflect RA-dependency rather than DV axis patterning *per se*, somite-derived signals in the human embryo could certainly contribute to DV axis patterning alongside the influence of the notochord. Previous research in animal models has suggested that RA signalling indeed plays an important role in DV patterning, but its precise function is often difficult to isolate due to the complex interactions with other signalling pathways, including Shh, Wnt, and FGF pathways. For instance, VAD quail embryos exhibit a ‘dorsal shift’ in neural tube patterning, with expanded domains of genes associated with a ventral identity compared to wildtype^80^, suggesting a dorsalising role for RA signalling. However, this interpretation is complicated by the concomitant increase in Shh expression in the neural tube, which is normally restricted to the floor plate. In such complex systems, it is difficult to delineate the respective roles of interacting pathways. Embryo models like hTLS might therefore allow us to isolate pathways, disentangling the roles of different signalling mechanisms in tissue patterning. In this study, we observed that <u>combined</u> exposure of neural tube cells to both SAG and somite-derived RA led to increased ventralisation, suggesting that the cells integrate the two pathways in a combinatorial manner. We show that even in the absence of a localized Shh source, RA signals emanating from the somites are sufficient to drive patterning in the neural tube. Further investigation into the molecular mechanisms underlying this signal integration, and how these pathways contribute to boundary formation during human neural tube DV patterning, could provide deeper insights into this interesting observation.

The multilineage composition of the hTLS model is a crucial feature, as it allows for the study of inter-tissue interactions, which are essential for proper embryonic patterning and organization. These interactions provide insights into the regulatory logic that governs multiscale coordination within the developing embryo, or co-development. However, while the hTLS model offers many advantages, it is important to recognize its limitations. For example, our data suggest that the notochord is not strictly required for neural tube-derived signalling to the somites, but we cannot exclude the possibility that additional relay interactions—such as notochord-induced floor plate maturation—may occur in the embryo, potentially influencing somite development. Similarly, while we observe that *ALDH1A2* expression is excluded from the lateral somites, we cannot determine whether this exclusion has functional implications for the development of the neighbouring intermediate or lateral plate mesoderm in the human embryo. These considerations highlight the distinction between embryo models and embryos, emphasizing the complementary role of such models in developmental research.

Moreover, we anticipate that hTLS models could be of value in disease modelling in the future, given their origin from human induced pluripotent stem cells. One notable observation from our study is the formation of the hTLS neural tube via a secondary neurulation-like process, which could prove valuable for investigating rare neural tube defects (NTDs). This is particularly interesting given that in the embryo secondary neurulation only occurs in the posterior-most neural tube and implies that the mode of neurulation can be experimentally decoupled from the AP identity. Furthermore, the ability to generate hTLSs from both ESCs and iPSCs with high reproducibility offers the potential for modelling patient-specific phenotypes, providing a platform for investigating disease mechanisms.

Overall, we have shown that interactions between the somites and neural tube leads to patterning events including mediolateral *ALDH1A2* expression in somites and elements of a DV-like axis in the neural tube, mediated through localised RA signalling. All of these events are spontaneous and self-organised, and can enable us to explore how feedback and relay mechanisms combine to enable co-development across tissues in a human developmental context.

## Methods

### Mouse work

All experiments carried out on mice were conducted according to the UK Animal (Scientific Procedures) Act under license PP6551133. The CD-1 (BRCD) animals were held within a Techniplast IVC green line system, and the air movement in the cages regulated by an Techniplast air management unit on negative pressure, 75 ACH/-20%, with all cages placed on automatic watering. The humidity and temperature were regulated at room level, set at code of practice standard levels of 20-24 °C and 55% (+/- 10%) humidity. The light cycle was 7am-7pm including dawn and dusk settings. The animals were kept on Datesand Eco Pure Chips sawdust, Bed-r’Nest nesting and smart homes enrichment, with Teklad Global Rodent Diet sterilised 2018S 18% Protein.

Embryos were obtained from pregnant mothers at E9.5 and dissected from maternal and extraembryonic tissues. Before fixation, embryos were washed in PBS for few minutes. Mouse embryos were fixed overnight in 4% paraformaldehyde (PFA; v/v) in RNase-free PBS (Invitrogen, AM9625) at 4 °C, then dehydrated gradually into methanol by incubating them for 5 min in a series of increasing concentrations (0%, 25%, 50%, 75% and 100% by volume, respectively) in RNase-free PBS with 0.1% (v/v) Tween20 (Sigma Aldrich, P1379) (PBST) on ice.

### Human Stem Cell lines

The cell lines used in this study include the human iPSC cell lines HES7::Achillies^48^, SOX2::mEGFP(FCI-AICS-0074-026)^86^, ZO-1::mEGFP (FCI-AICS- 0023)^86^, β-catenin::mEGFP (FCI-AICS-0058 cl.67)^86^, LaminB1::mTagRFP-T (FCI- AICS-0034 cl.62)^86^ and H2B::mEGFP (FCI-AICS-0061 cl.36)^86^; and the human ES cell line RUES2-GLR^46^. This research used HESCU core facility expanded cell lines made possible through the Allen Cell Collection, available from Coriell Institute for Medical Research. All cells were cultured in incubators at 37 °C and 5% CO_2_. Human iPS cells were cultured routinely in either StemFit Basic 02 (Ajinomoto, discontinued in EU) supplemented with 80 ng/mL bFGF (Peprotech, 100-18B) or mTeSR^TM^ 1 (STEMCELL Technologies, 85850) on 0.25 µg/cm^2^ recombinant Laminin iMatrix-511 silk E8- coated plates (AMSbio, AMS892 021). Cultures of 80-90% confluency were routinely passaged using Accutase^TM^ (STEMCELL Technologies, 07922) and single cells were seeded with medium supplemented with 10 µM Y- 27632 (ROCK inhibitor; Tocris Biosciences, 1254). Fresh medium was exchanged daily. Cells were regularly tested for Mycoplasma contamination by PCR.

### Human Gastruloids

RUES2-GLR hESCs grown routinely in NutriStem^®^ hPSC XF were first plated as single cells at a density of 5.2-6.7 x 10^3^ cells/cm^2^ with 5 µM Y-27632 (Tocris Biosciences, 1254) for the first 24h after plating. After further culture, they were pretreated on the fourth day (96h) with 4 µM CHIR99021 (Chiron; Tocris Biosciences, 4423) and 10 µM SB431542 (Tocris Biosciences, 1614; TGFβ inhibitor) in NutriStem^®^ hPSC XF for 24 hours. After this, they were dissociated in StemPro™ Accutase™ (Gibco™ Thermo Fisher Scientific, A1110501) and 700 cells were seeded into each well of an ultra-low-adherence 96-well plate (Greiner Bio-One, 650970) in 50 μl Essential 6 medium (Gibco™ Thermo Fisher Scientific, A1516401) supplemented with 5 µM Y-27632 and 3 µM Chiron. Medium was replaced daily with fresh Essential 6. At 76 hours, the structures were embedded in 20% (v/v) Corning^®^ Matrigel^®^ growth factor-reduced basement membrane matrix, phenol red-free, LDEV-free (Corning, 356231) or Corning^®^ Matrigel^®^ hESC-qualified matrix, LDEV-free (Corning, 354277); in Essential 6 medium to a final protein concentration of 1.96 mg/mL. The 20% (v/v) Matrigel solutions were prepared on ice with ice-cold Essential 6, before plating as 5 μL droplets onto tissue-culture treated plastic on ice.

Human gastruloids were collected and transferred individually into separate 5 μL droplets with a micropipette. After transfer, droplets were left to gelate for 1h in the incubator at 37°C and 5% CO_2_, before covering gently with excess Essential 6 medium pre-warmed to 37°C.

### Generating human trunk like-structures

Once the human iPSCs were around 70-80% confluency, cells were dissociated in Accutase^TM^ and seeded at a density of 20,000 cells per well of a Laminin-coated 6-well plate with 2 mL StemFit 02 with 80 ng/mL bFGF supplemented with 10 µM Y-27632. After 5 days, cultures were pre-treated in StemFit 02 with 20 ng/mL bFGF supplemented with CHIR99021 (Chiron; Tocris Biosciences, 4423). Alternatively, cultures maintained in mTeSR1 were pre-treated with N2B27 supplemented with CHIR99021. N2B27 medium was prepared from 1:1 mix of Advanced DMEM/F12 (Gibco, 12491015) and Neurobasal medium (Gibco, 21103049) supplemented with N-2 Supplement (100×) (Gibco, 17502048), B27 supplement (50×) without Vitamin A (Gibco, 12587010) and 1% GlutaMAX Supplement (v/v) (Gibco, 35050061). The optimal concentration of Chiron for pre-treatment needed to be adjusted depending on the cell line and medium used but ranged between 2-5 µM. Concentrations for the lines used in this study are described in Table 1.

After 24 h of pre-treatment, cells were dissociated with Accutase^TM^. Cell numbers were determined using an automated cell counter (NuceloCounter® NC- 202^TM^, ChemoMetec) and 500 cells were seeded into one well of an ultra-low-adherence 96-well plate (Greiner Bio-One, 650970) with 50 μL either StemFit medium with 20 ng/mL bFGF or N2B27 (depending on the medium used for pre-treatment) and supplemented with 10 µM Y-27632, 10 µM SB431542 (TGFβ inhibitor) and cell-line dependent concentration of Chiron (Table 1). The plate was centrifuged for 160 *g* at room temperature for 2 min to form a cell pellet at the bottom of the well. The following day, 150 μL fresh StemFit 02 or N2B27 was added to each well and exchanged again after 24 h with 150 μL of medium. At 72h after aggregation, 150 μl of medium was removed from aggregates and they were embedded into medium containing 10% (v/v) growth factor-reduced Matrigel (Corning, 356231) and 1 µM all *trans*-retinal (Tocris, 0695), unless otherwise stated. No further medium changes were required from this point.

### Scanning Electron Microscopy

Human trunk like-structures, made from LaminB1-RFP line at 120h after aggregation, were recovered from Matrigel using Cell Recovery Solution (Corning, 354253) for 30 mins at 4 °C, washed twice with Dulbecco’s phosphate-buffered saline without Mg^2+^ or Ca^2+^ (PBS; Gibco, 14190144) and fixed overnight in 4% PFA (v/v) in PBS (PFA; Alfa Aesar, 43368.9M) at 4 °C.

They were then additionally fixed in 4% PFA/ 2.5% glutaraldehyde (v/v) in 0.1M phosphate buffer (PB) for 1 hour at room temperature (RT), washed 3x for 5 minutes in 1% bovine serum albumin (BSA, v/v) in 0.1M PB, postfixed in 1% reduced osmium (1% OsO_4_/ 1.5% K_3_Fe(CN)_6_; v/v) for 1 hour at 4°C, washed 3x 5mins in PB, and then dehydrated with a graded ethanol series (70%, 90% and 2 x 100% by volume) for 20mins each at RT. The samples were then critical point dried (Leica EM CPD300 Critical Point Dryer), mounted on stubs, sputter coated with 5 nm of platinum (Quorum 150RS) and then imaged on the Quanta 250 FEG SEM (Thermo Fisher Scientific).

The SEM was operated between a chamber pressure of 4.0-4.38 Pa with 30 μm aperture inserted using an accelerating voltage of 2.5kV (Figure 1e – right, top) 3 kV (Figure 1e - left) and 5kV (Figure 1e – right, middle and bottom), the dwell time 20 μs and a working distance of 7.4 mm (Figure 1e – right, bottom), 7.6 mm (Figure 1e – right, middle), 9.2 mm (Figure 1e - left) and 11.6 mm (Figure 1e – right, top). A 90° tilt sample holder was used for Figure 1e, right, top panel.

### Immunostaining

Pre-embedded structures were transferred into 1.5 ml tubes for fixation. Some structures were first recovered from Matrigel using Cell Recovery Solution for 30 mins at 4 °C (Figure 4e,g; Figure 5c,f,g; Supplementary Figure 8d; Supplementary Figure 9d,e), although we noticed some lack of morphological integrity upon exposure so this was not used routinely. They were then washed with PBS twice, fixed overnight in 4% PFA (v/v) at 4 °C, and then washed again 3 times for 10 mins in PBS. Then, samples were washed 3 times for 10 min each in PBS with 0.2% Triton X-100 (v/v) and 3% BSA (w/v, PBS-BT) at room temperature and blocked for 2 hours in PBS-BT at 4 °C. Primary antibodies were diluted in blocking solution and incubated overnight at 4 °C, with gentle rocking. Following several washes in PBS-BT (2 x 5 min at room temp; 3 x 10 min at 4 °C; 4 x 30 min at 4 °C), samples were incubated with Alex-Fluor-conjugated secondary antibodies with 1:500 Phalloidin CruzFluor 647 (Santa Cruz, sc-363797) or 1:800 Hoechst 33342 (Invitrogen, H3570) in PBS-BT with 10% normal donkey serum (v/v) overnight at 4 °C with gentle rocking away from light. After additional washes (3 x 10 min in PBS-BT at room temp, 2 x 5 mins in PBS with 0.2% Triton X-100 (Merck, T8787) and 0.2% BSA (PBT) and 3 x 15 mins in PBT at room temp), samples were mounted in Vectashield (Vector Labs, H- 1900-10) onto either a Superfrost microscopy slide (Epredia, AA00008032E01MNZ10) or coverslip (VWR, 631-0138).

The antibodies used in this study were: 1:200 rabbit anti-Brachyury (Abcam, ab209665); 1:500 mouse anti-MEOX1 (ThermoFisher, TA-804716); 1:200 goat anti- SOX2 (R&D Systems, AF2018-SP), 1:400 rabbit anti-PAX6 (ThermoFisher, 42-6600) and 1:10 mouse anti-PAX7 (DSHB, PAX7). All secondary antibodies were raised in Donkey and diluted 1:500, and included Alexa Fluor 488-, 594- and 647 conjugated antibodies (Jackson ImmunoResearch).

### Hybridisation Chain Reaction

Custom probe sets (all probes set size = 20 pairs), HCR amplifiers and buffers (Hybridisation, wash and amplification buffers) were purchased from Molecular Instruments. Pre-embedded structures were fixed in 4% PFA (v/v) in RNase-free PBS overnight as described above. Samples were dehydrated gradually into methanol by incubating them for 5 min in a series of increasing concentrations (0%, 25%, 50%, 75% and 100% by volume, respectively) in PBST and stored at -20°C. For HCR, samples were rehydrated gradually by incubating them for 5 mins in a series of decreasing concentrations of methanol (75%, 50%, 25% and 0% by volume, respectively) in PBST. Samples were washed once more in PBST, then incubated for 5 mins with hybridisation buffer at room temperature (RT), then for a further 30 min at 37 °C, alongside a combination of probes (8 nM final) in hybridisation buffer. Samples were then incubated with the probe solution overnight at 37 °C. To remove excess probes, samples were washed with probe wash buffer 4 times for 15 min each at 37 °C, followed by a further 3 washes in 5× SSCT 3 times for 5 min at RT. Samples were pre-amplified in amplification buffer for 30 min at RT, gently rocking. Amplifier hairpins h1 and h2 were snap-cooled separately by first heating at 95 °C for 90 s and then cooling down at RT for 30 min in the dark. Hairpin amplifiers were diluted to 6 nM in amplification buffer, added to samples and then incubated overnight in the dark at RT. Next, samples were washed in 5× SSCT 4 times for 5 min, then 2 times for 30 min and with a final wash in PBST for 2 min. Samples were counterstained 1:1000 with Hoechst 33342 overnight at 4 °C. Next day, samples were washed 3 times with PBST for 5 mins and postfixed with 4% PFA (v/v) in RNase-free PBS for 20 mins and then washed a further 3 times with PBST for 5 mins at 4 °C with gentle rocking. Samples were mounted in Vectashield or Prolong Glass Antifade Mountant (Invitrogen, P36980) onto either a Superfrost microscopy slide or coverslip, then cover-slipped. HCR probes and the associated amplifiers used are listed in Table 2.

For HCR of mouse embryos, samples were rehydrated gradually by incubating them for 5 mins in a series of decreasing concentrations of methanol (75%, 50%, 25% and 0% by volume, respectively) in PBST. Embryos were bleached in 6% H_2_O_2_ (v/v) in PBS for 15 mins and were washed 3 times for 5 min each in PBST at room temperature. Samples were treated with 10 μg/mL Proteinase K (Roche, 3115879001) for 10 mins (no rocking) and then washed twice for 5 mins in PBST (no rocking). Afterwards, embryos were post-fixed for 20 mins with 4% PFA (v/v) in RNase-free PBS at room temperature, gently rocking and then washed 3 times for 5 mins with PBST. To equilibrate embryos, samples were washed in 1:1 prewarmed Hybridisation Buffer and PBST for 5 mins. Embryos were washed with prewarmed Hybridisation Buffer for 5 mins before following the HCR protocol as described above.

### Imaging

#### Confocal Imaging

Confocal imaging was performed using a Zeiss Invert880 with Airyscan 2014 or Zeiss Invert880 NLO OPO, using a Zeiss Pan-Apochromat 20x/0.8 objective. Z-stacks were obtained with 5 µm steps, averaging each slice 4 times, unless otherwise stated. Any tiled images were acquired with a 10% overlap. Spectral unmixing was used for imaging HCR samples, unless otherwise stated (Track method used for Figure 1d, Figure 2e, Figure 3h, Figure 5d-e and Supplementary Figure 1i). Briefly, spectra were set up for all HCR amplifiers by imaging droplets of snap-cooled mixed hairpin pairs using lambda mode and then areas selected to save each spectrum to a spectral database. HCR images were imaged with lambda mode and linear unmixing was run with saved spectra of amplifiers, nuclear signal and autofluorescence spectra of hTLS. Confocal images are presented either as optical slices or maximum intensity projections, generated using FIJI or Imaris (Oxford Instruments), unless otherwise stated. For Figure 5g, XY images and YX optical slices were taken of hTLS imaged in Figure 5e using average mode in Imaris.

#### Live Imaging

Brightfield widefield or confocal spinning disk images on live samples were obtained on ImageXpress Confocal HT.ai system with Nikon 10X S Plan Apo Lambda objective. Time-lapse imaging was acquired with intervals between 20-30 minutes under environmental control at 37°C and 5% CO_2_.

#### Lightsheet Imaging

After post fixation following the HCR protocol, mouse embryos for lightsheet imaging were embedded in a cryomold in warm 1% Ultralow melting agarose (Sigma, A5030) and placed on ice to polymerise. Samples were then prepared for clearing with BABB. All steps were carried out in a glass vial, which was covered with foil. Mouse embryos were dehydrated gradually into methanol by incubating them for 1 hour at room temperature with gentle rocking in a series of increasing concentrations (50%, 70%, 80%, 90%, and twice in 100% by volume, respectively) in deionised water. Embryos were then incubated in a solution of 1:1 100% methanol and BABB (1:2 solution of Benzyl alcohol (BA; Sigma, 24122) and Benzyl benzoate (BB; Thermo Fisher, 105860010)) for 1 hour at room temperature with gentle rocking. Samples were then transferred into 100% solution of BABB and left overnight. Next day, BABB was removed, and samples were washed in Ethyl cinnamate (ECi; Sigma 112372) for 5 mins before transferring samples into fresh ECi for storage. Mouse embryos were imaged using Miltenyi-Lavision UM II with ECi as the imaging solution.

Z-stacks were obtained with 4 µm steps with dynamic focus. Lightsheet images and videos were generated with Imaris.

#### Image Analysis

*NMP Colocalisation Analysis (Whole Volume Segmentation):* Confocal images of hTLS stained with Hoechst, TBXT, and SOX2 were processed through Imaris. Surfaces of whole hTLS were generated from the Hoechst, TBXT, and SOX2 channels. A new channel of the colocalised region was generated using the colocalisation package in Imaris and then a new surface was generated from this channel. Total volume for each channel was then measured.

*Somite polarisation of gene expression:* Confocal images were imported into Imaris and the Hoechst channel used to create a manual surface rendering of the whole hTLS, the individual somites and the neural tube. New channels were created for fluorescent *UNCX* and *ALDH1A2* HCR signal within each somite, and this was then used to generate surfaces corresponding to positive signal within each somite.

*Morphometric analysis:* The pixel classification tool Ilastik^87^ was used to label hTLS (24h-144h) in transmitted light images to generate probability maps, distinguishing the hTLS from background. CellProfiler^88^ was used to obtain basic morphometry through the generation and measurement of objects using these probability maps. Objects touching the edge of image and those below a pixel threshold were automatically discarded. Images were then manually checked, and sample images containing fibres and debris discarded. Medial length was obtained through CellProfiler via ImageJ^89^: objects were skeletonised, and the largest, shortest branch length selected. Quantification of somite number was measured through manual counting of segmented structures on 96-144h hTLS.

*Oscillation quantification:* HES7-Achilles samples were imaged on the ImageXpress Confocal HT.ai system from 100-122 hours in the hTLS protocol. Initial image analysis was performed in FIJI^90^ where ROIs for HES7 oscillations and background were selected. Integrated density, mean pixel intensity and area were measured and inputted into the following formula: Corrected Total Fluorescence = Integrated Density – (Area of selected cell X Mean fluorescence of background readings). Detrending was then performed in terminal on MacOS, and peaks/valleys were calculated in Excel and plotted in RStudio.

#### Signalling manipulation

For RA signalling manipulation of hTLS, retinal (RAL) was replaced with either DMSO or all *trans*-retinoic acid (100 nM; Sigma Aldrich, R2625); or treated with BMS493 (2.5 µM; Tocris 3509) at 72h during embedding. Additionally, for 96h RA inhibition of hTLS, BMS493 was added to 20 µL of medium containing 10% growth factor-reduced Matrigel (v/v) and 1 µM all *trans*-retinal and gently added to each well for a final concentration of 2.5 µM. For Shh signalling activation of hTLS models, SAG (Tocris, 4366) was added to 20 µL of medium containing 10% growth factor- reduced Matrigel (v/v) and 1 µM all *trans*-retinal and gently added to each well for the following final concentrations 100 nM, 200 nM, 300 nM, 400 nM and 500 nM.

#### Single Cell RNA-Sequencing

Samples from each time point were collected from 96WPs and pooled together. TrypLE Select was added to the samples and incubated at 37°C for 15-20 minutes. Samples were gently pipetted up and down until hTLS had been dissociated into single cells. The cells were then spun in the centrifuge at 170 x g for 4 minutes and resuspended in PBS^-/-^ + 0.04% BSA (v/v) and kept on ice until further processing. The concentration and viability of the single cell suspension was measured using acridine orange (AO) and propidium iodide (PI) with a Luna-FX7 Automatic Cell Counter. 13200 cells were loaded onto a Chromium Chip and partitioned in nanolitre scale droplets using the Chromium Controller and Chromium Next GEM Single Cell Reagents (CG000315 Chromium Single Cell 3’ Reagent Kits User Guide (v3.1 - Dual Index)). Within each droplet the cells were lysed, and the RNA was reverse transcribed. All of the resulting cDNA within a droplet shared the same cell barcode. Illumina-compatible libraries were generated from the cDNA using Chromium Next GEM Single Cell library reagents in accordance with the manufacturer’s instructions (10x Genomics, CG000315 Chromium Single Cell 3’ Reagent Kits User Guide (v3.1 - Dual Index)). Final libraries are Quality Controlled using the Agilent TapeStation and sequenced using the Illumina NovaSeq 6000. Sequencing read configuration: 28-10- 10-90.

#### Bioinformatic Analysis

FASTQ files were aligned to the hg38 transcriptome, and count matrices were generated, filtering for GEM cell barcodes (excluding GEMs with free-floating mRNA from lysed or dead cells) using CellRanger (version 6.0.1). Count matrices were imported into R (version 4.1.0) and processed using the Seurat library (version 4.4.0) following the standard pipeline^91^. Low-quality cells were removed, with cells kept for further analysis if they met the following criteria: the mitochondrial content was within three standard deviations from the median, more than 500 genes were detected and more than 1,000 RNA molecules were detected. DoubletFinder was used to identify doublets, assuming a theoretical doublet rate of 7.5%, which were removed from subsequent analysis^92^.

All samples were integrated using the RPCA method, implemented by Seurat’s functions FindIntegrationAnchors() and IntegrateData(), using the top 3,000 variable features and the first 25 principal components as determined using the ’maxLikGlobalDimEst’ function from the intrinsicDimension package (version 1.2.0). Effects of cell cycle heterogeneity was calculated using cell cycle phase scores based on canonical markers and regressed out of the data using ScaleData. Eight clusters were identified at a resolution of 0.5.

For RNA velocity analysis, loom files containing spliced and unspliced matrices were calculated, from BAM files generated by CellRanger, using the package velocyto (version 0.17.8) and Python 3.6.4. The matrices were used as input to scVelo (version 0.2.4) to calculate the RNA velocity values for each gene of each cell using the package loompy (verions 3.0.6) and scanpy (version 1.8.2). scVelo was used in the dynamical mode with default settings. The resulting RNAvelocity vector was overlaid onto the UMAP space by translating the RNAvelocities into likely cell transitions using cosine correlation to compute the probabilities of one cell transitioning into another cell.

Lineages for Trajectory analyses were identified for the Somite and Neural fates separately using the NMP cluster as the starting population using Monocle3 (version 1.3.0)^93^. Differentially expressed genes along each lineages were identified with the ’graph_test’ function using the Morans I test. The top 100 genes were used to draw smoothed heatmaps using Tradeseq (version 1.7.07)^94^ and ComplexHeatmap (version 2.2.0)^95^.

Cell-to-cell crosstalk was inferred using the R library CellChat (version 2.1.2)^71^ and the CellChat human database. Clusters designated NP1, NP2, NP3 and Neuronal were merged to create one cluster. Only the Somites and the merged Neuronal clusters were considered for Cellchat analysis. The CellChat analysis was performed as outlined in the CellChat vignette, with the ’population.size’ parameter set to TRUE when computing the communication probability between clusters. Significant interactions were filtered on inter-cluster and intra-cluster interactions were removed.

Published datasets were reprocessed using Seurat^91^ v4.4.0 from either the counts matrix of a Seurat object or downloaded outputs of CellRanger from NCBI’s GEO repository.

Four samples (comprising of hAxioloid_48h, hAxioloid_72h, 96h_MG_RAL and 120h_MG_RAL from the axioloid dataset (GSE199576; ^15^) were downloaded as count matrixes and processed using Seurat version 4.4.0. Samples were integrated using methods highlighted in the authors publication and cells assigned clusters according to published cell metadata. Cells from this dataset were projected onto our integrated dataset using the functions ‘FindTransferAnchors’ (using the first 30 dimensions of the pca space) and ’MapQuery’. To determine similarity between axioloid clusters and hTLS integrated clusters, a Spearman’s rank correlation was calculated on the aggregated cell expression using the mean gene expression per cluster and per sample, after filtering on common genes between datasets.

The full human embryo dataset (GSE157329; ^37^] was downloaded and processed using scripts provided by the authors (https://github.com/zhirongbaolab/heoa) with Seurat 4.4.0. Cells were annotated using published annotation. Predicted cell type cluster and stage annotation scores were filtered using a threshold of > 0.5.

The human neural tube dataset^68^ was used to look for similarity of clusters within our integrated Neuronal clusters (labelled NP1, NP2, NP3 and Neuronal). A Spearman’s rank correlation test was used on aggregated cell expression per cluster using common genes between both datasets.

### Ethical Statement

The hTLS model as described in this manuscript represents a human stem cell-based embryo model of post-gastrulation stages. However, the model lacks extraembryonic tissues (and is therefore ‘non-integrated’ by current ISSCR terminology^96^), as well as lacking tissues including anterior neural, intermediate- and lateral- mesoderm, notochord, endoderm, non-neural ectoderm amongst others. As such, we believe the system represents a minimal developmental model.

### Data Availability

The sequencing datasets generated during the current study are available in the GEO repository, GSE268451 with accession codes GSM8291523-5 (DOI link will be provided on publication).

## Acknowledgements

We would like to thank all those who helped and assisted with technical expertise on this project, including Matt Renshaw with confocal microscopy, Alessandro Ciccarelli with lightsheet microscopy, Aaron Sait and Lucy Collinson with scanning electron microscopy, Lyn Healy with human pluripotent cell line acquisition and culture guidance, Emma Connick and Robert Gunn with single cell RNA sequencing. In addition, we are grateful to Ali Brivanlou, Olivier Pourquie and the Allen Cell Collection (through the Coriell Institute for Medical Research) for kindly sharing their cell lines. We also thank James Briscoe and Alfonso Martinez Arias as well as members of the Developmental Models lab for their feedback on the manuscript.

This work was funded by an MRC-AMED grant (MR/V005367/2) as well as funding from the Francis Crick Institute which receives its core funding from Cancer Research UK (CC2186), the UK Medical Research Council (CC2186) and the Wellcome Trust (CC2186).

## Author Contributions

K.M.: Methodology, Formal Analysis, Investigation, Data Curation, Writing - Review & Editing, Visualization. L.T.: Methodology, Software, Formal Analysis, Investigation, Data Curation, Writing - Review & Editing, Visualization. P.C.: Methodology, Software, Formal Analysis, Resources, Data Curation. J.T.: Methodology, Formal Analysis, Investigation, Data Curation. I.R-P.: Investigation, Writing - Review & Editing. P.B-B. Investigation, Writing - Review & Editing. N.M.: Conceptualisation, Methodology, Writing - Original Draft, Writing - Review & Editing, Visualization, Supervision, Project administration, Funding acquisition.

