## Supplementary Information for "Modelling co-development between the somites and neural tube in human Trunk-like Structures (hTLS)"

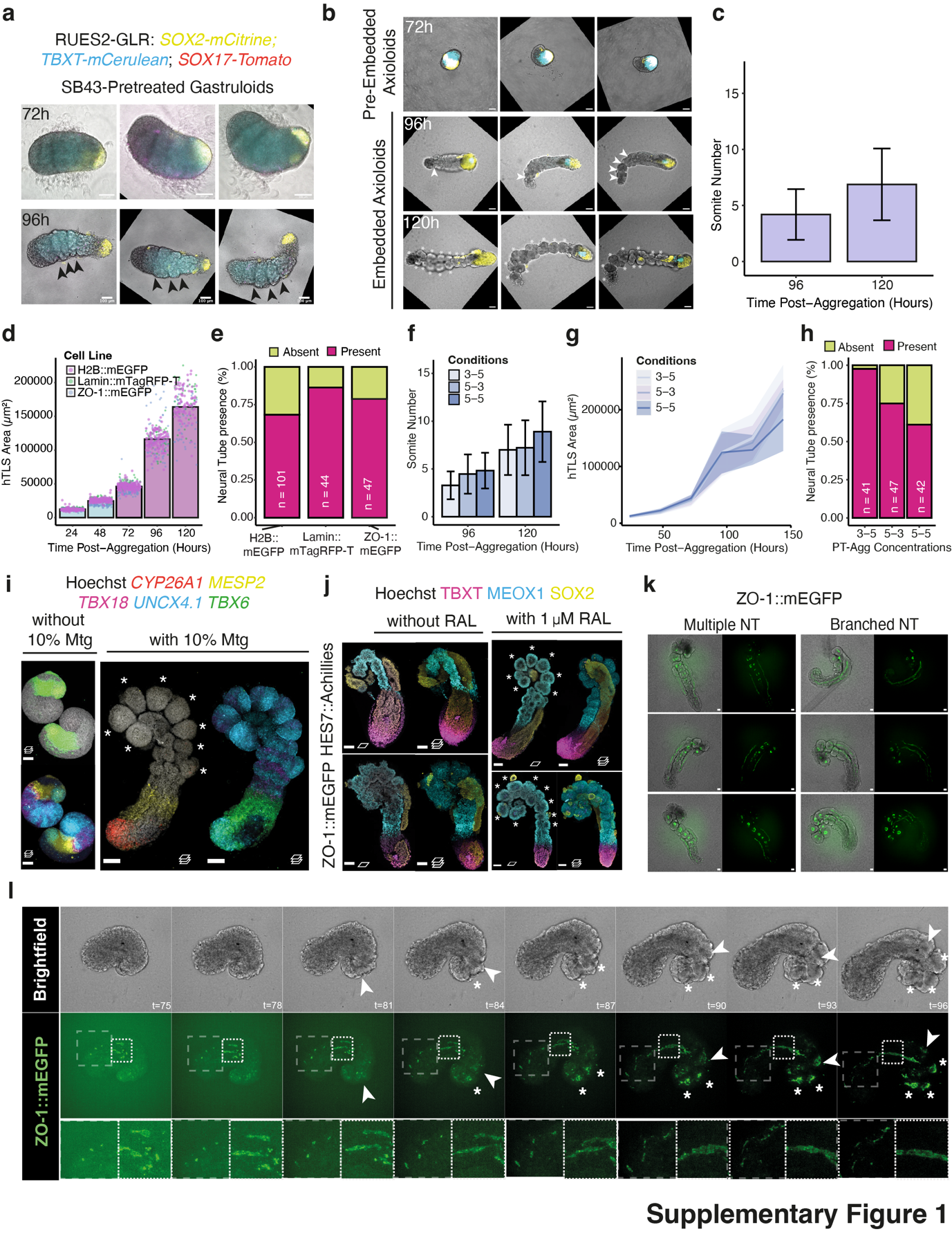

**Supplementary Figure 1: Establishing and characterising the hTLS model. (a)** Human gastruloids pretreated with the Nodal signalling inhibitor, SB43 (SB431542), show a typical ‘striped’ pattern at 72h (top), and sequential somite-like structures upon embedding in Matrigel (bottom). Black arrowheads indicate somite-like structure boundaries. **(b)** Morphology of RUES2-GLR axioloids over time. Three representative images are shown. Scale bars, 100 µm. White arrowheads indicate segmenting boundaries, * indicates individual segments. **(c)** Number of somites in ZO-1::eGFP hTLS over time (mean±standard deviation(sd)). **(d)** Change in size of ZO-1::GFP, LaminB1::RFP, and H2B::GFP hTLSs over time (bar indicates mean average). **(e)** Neural tube presence in 120h hTLS across cell lines. **(f-h)** hTLS morphology quantifications following varied exposure to Chi (CHIR99021) at pretreatment (PT) and aggregation (Agg) stages. Cultures were pre-treated with 3 µM Chi, and aggregated with 5 µM Chi (‘3-5’), pre-treated with 5 µM Chi, and aggregated with 3 µM Chi (‘5-3’), or pre-treated with 5 µM Chi, and aggregated with 5 µM Chi (‘5-5’). **(f)** Change in somite number over time. **(g)** Change in hTLS size across time. **(h)** Proportion of structures with neural tube present or absent. (**i)** Projected images of 120h hTLS with and without embedding in Matrigel (Mtg). Scale bars, 100 μm. * indicates somites. **(j)** Immunostained optical slice (left) and projections (right) of 120h hTLS with and without exposure to retinal (RAL) from 72-120h. These hTLS were generated from HES7::Achillies (top) and ZO-1::eGFP (bottom) cell lines. Scale bars, 100 μm. * indicates somites. **(k)** Representative images of ZO-1:eGFP hTLSs, showing multiple and branched neural tubes (NT). Scale bars, 50 μm. **(l)** Representative brightfield (top) and confocal images (bottom) of live imaging of ZO-1:eGFP hTLS at 3 hour intervals. White arrows indicate segmenting somite boundaries and * indicates formed somites. Dashed boxes highlight enlarged regions. Scale bars, 50 μm.

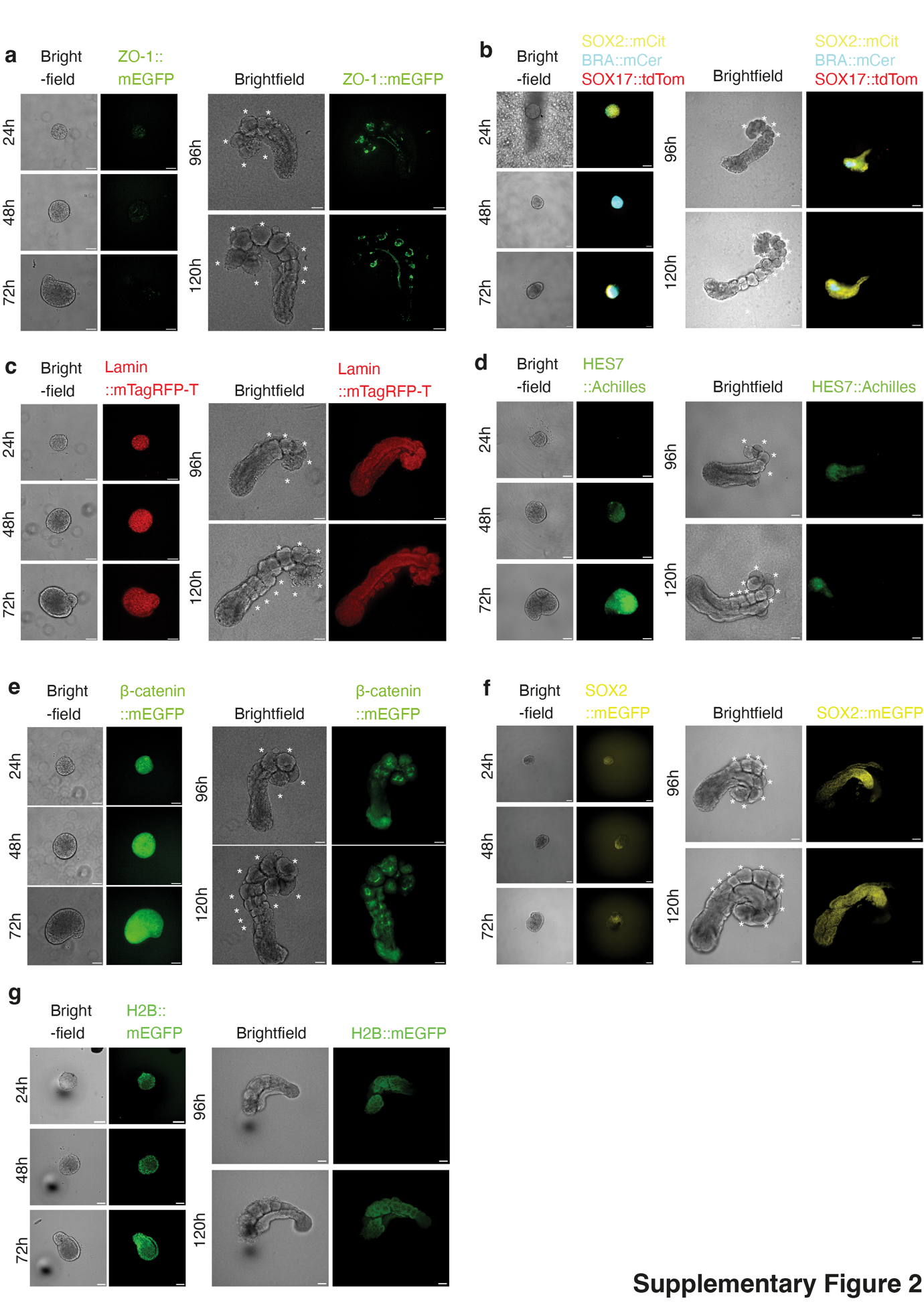

**Supplementary Figure 2: Generation of hTLS from several cell lines. (a-f)** Representative brightfield images of hTLS across timepoints, alongside fluorescent reporter expression for (**a)** ZO-1::eGFP, (**b)** RUES2-GLR, (**c)** LaminB1::RFP, (**d)** HES7-Achilles, (**e)** β-catenin::eGFP, (**f)** SOX2::GFP and (**g**) H2B::eGFP. Scale bars, 100 μm.

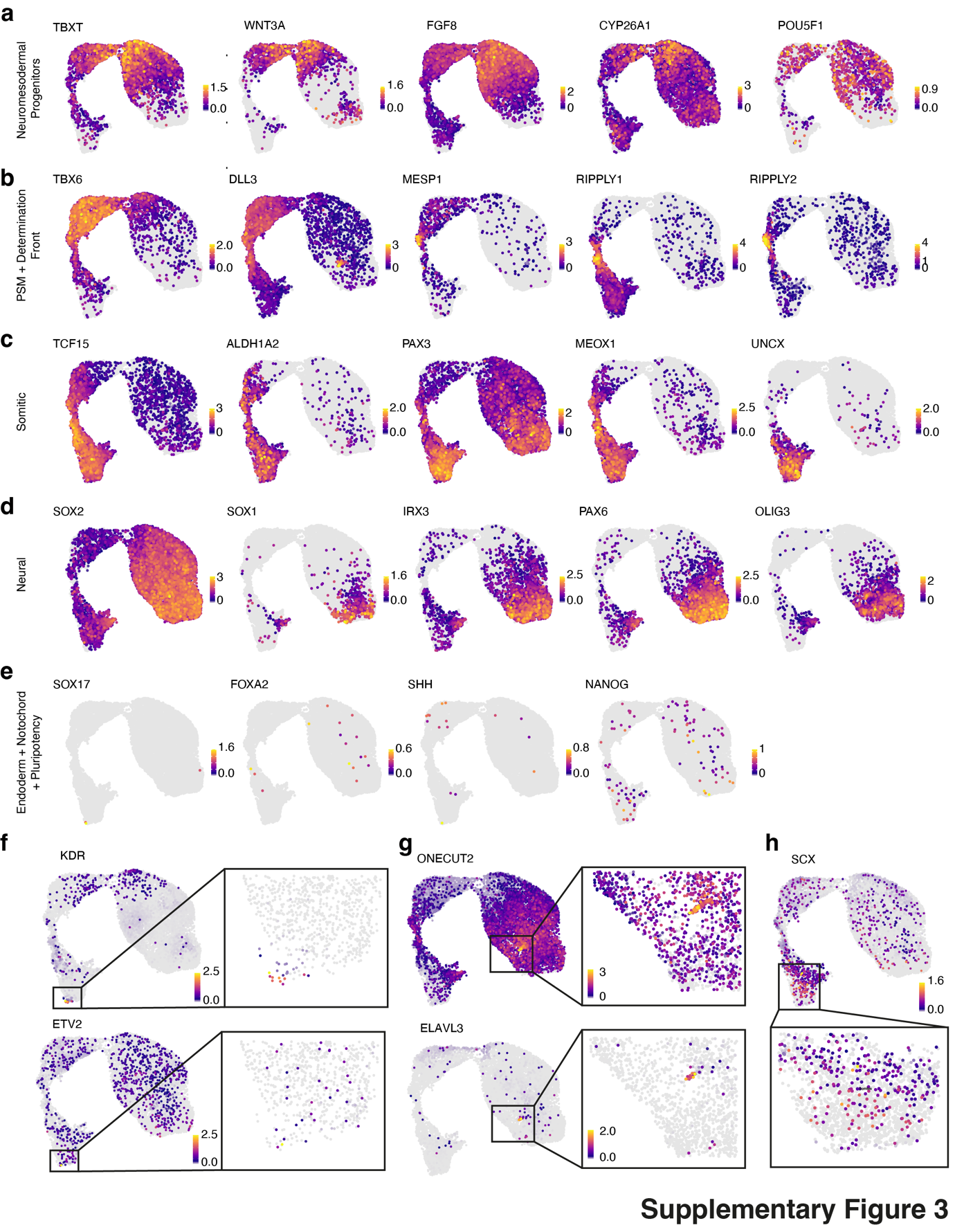

**Supplementary Figure 3:** **Gene expression signature of hTLS single cells.** Cells in UMAP space coloured by gene expression for markers of neuromesodermal progenitors **(a),** the presomitic mesoderm (PSM) and determination front (**b**), somitic mesoderm (**c)**, neural tissue (**d**), and markers of endoderm (*SOX17, FOXA2*), notochord (*SHH*) and pluripotency (*NANOG*; **e**). Zoomed-in regions are also shown for small sub-populations of cells expressing endothelial markers (KDR, ETV2; **f**), neural differentiation markers (ONECUT2, ELAVL3; **g**) and sclerotome (SCX, **h**).

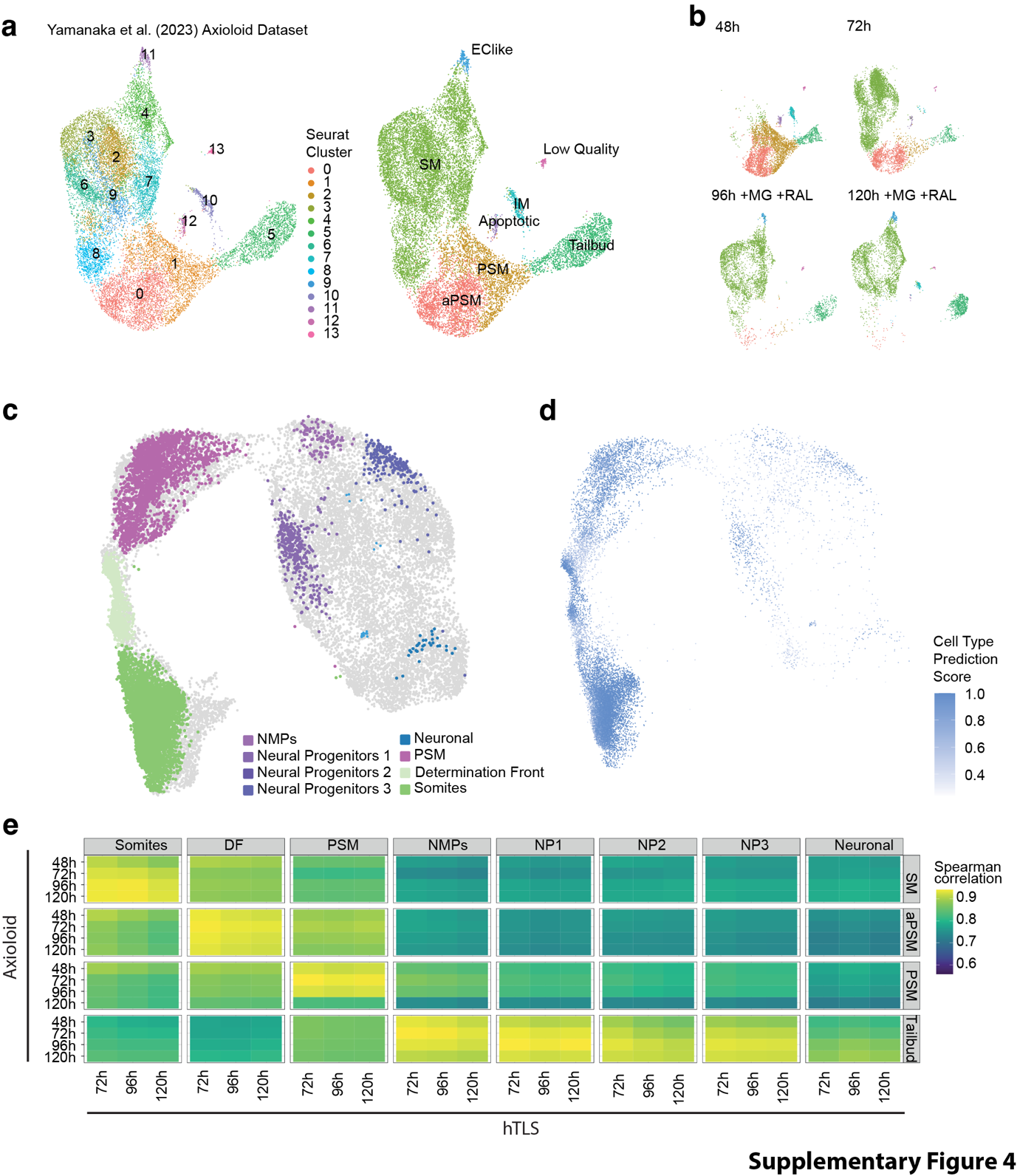

**Supplementary Figure 4: Comparison between hTLS and an axioloid dataset.** (**a**) UMAP projection of an axioloid dataset^15^, with clusters as identified by Seurat (left), alongside annotated map determined by marker gene expression and comparison to published labels. Clusters 2-4 and 6-9 were combined to form a larger cluster named ‘somitic mesoderm’ (right). (**b**) Position of cells in UMAP space according to their original sample across timepoints. (**c**) Projection of axioloid single cells onto the hTLS UMAP, coloured by label transfer from hTLS cluster identity. Note that the somitic lineages are well populated by axioloid cells, while the neural identies are mostly absent. (**d**) Projected cell coloured by prediction score as a measure of confidence. (**e**) Correlation in gene expression between timepoints in the axioloid and hTLS datasets, by cell cluster. IM, Intermediate mesoderm; EC, SM, Somitic mesoderm; PSM, Presomitic mesoderm; aPSM, anterior PSM; DF, Determination Front; NMPs, Neuromesodermal progenitors; NP, neural progenitor; MG, Matrigel; RAL, Retinal.

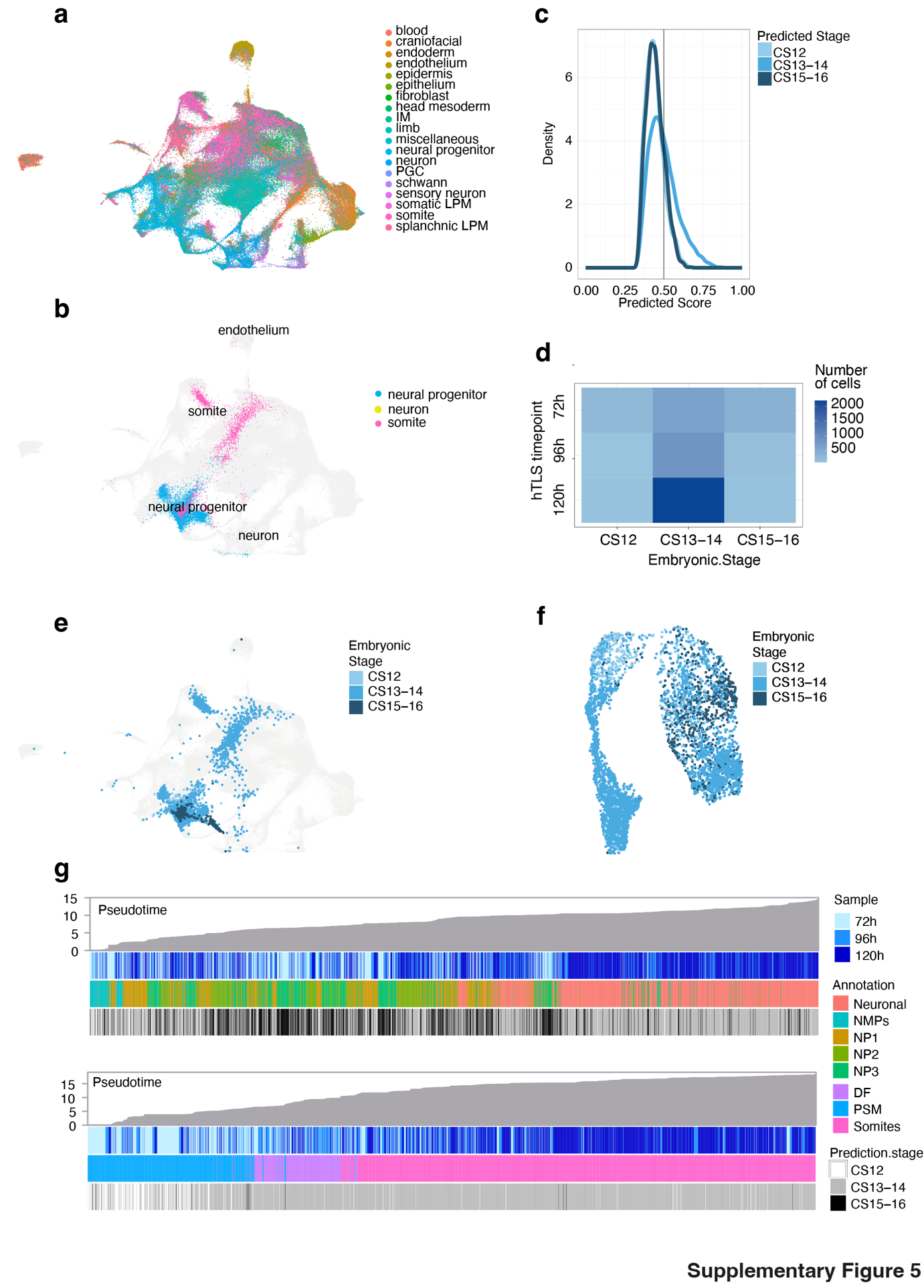

**Supplementary Figure 5: Bioinformatic comparison to a human embryo dataset.** (**a**) UMAP projection of a human embryo atlas^37^. (**b**) Projection of hTLS single cell data onto embryo atlas, and coloured according to label transfer (pink, somite; blue, neural progenitor; yellow, neuron). (**c**) Density plot of the predicted confidence score for each projected cell. Vertical line shows the cutoff used for label transfer and subsequent analysis = 0.5. (**d**) Number of hTLS cells assigned to human embryonic stages by label transfer, across the analysed timepoints (72-120h). (**e**) UMAP projection of hTLS cells onto human embryo atlas, coloured by equivalent embryonic stage according to label transfer. (**f**) UMAP projection of hTLS cells coloured by equivalent embryonic stage according to label transfer. (**g**) hTLS single cells ordered by pseudotime, showing sample, annotation and label transfer of equivalent embryonic stage. CS, Carnegie Stage.

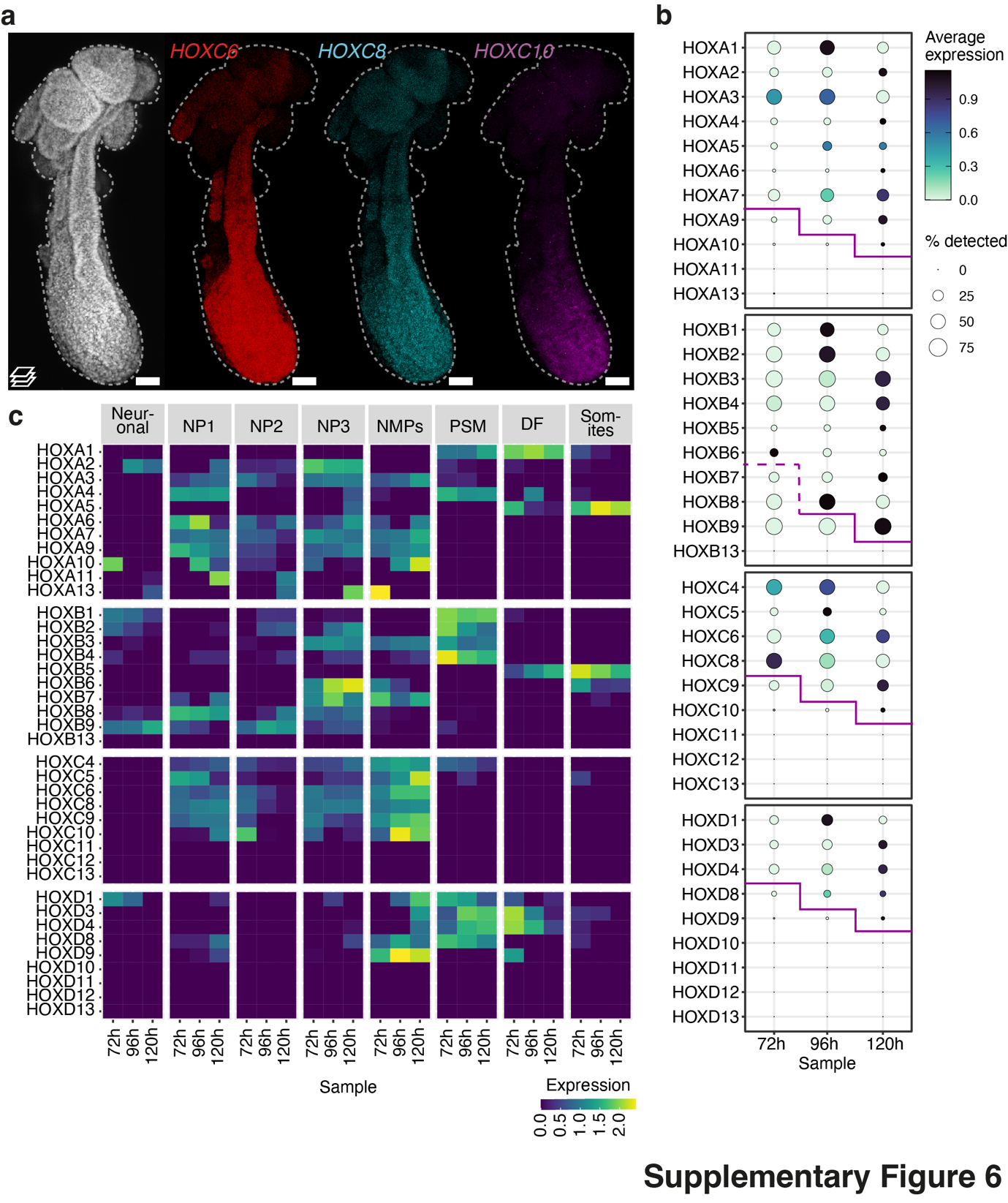

**Supplementary Figure 6: Spatiotemporal HOX gene expression. (a)** Projected HCR images of whole mount 120h hTLS. Scale bars, 50 μm. Dashed line outlines hTLS. (**b**) Temporal change in HOX gene expression from single cell transcriptomics. Purple line indicates the approximate maximal expression of HOX gene paralogues at each timepoint, dotted line indicates uncertainty in the HOXB cluster at 72h timepoint due to low frequency of detection. (**c**) Spatiotemporal gene expression of HOX genes across single cell clusters.

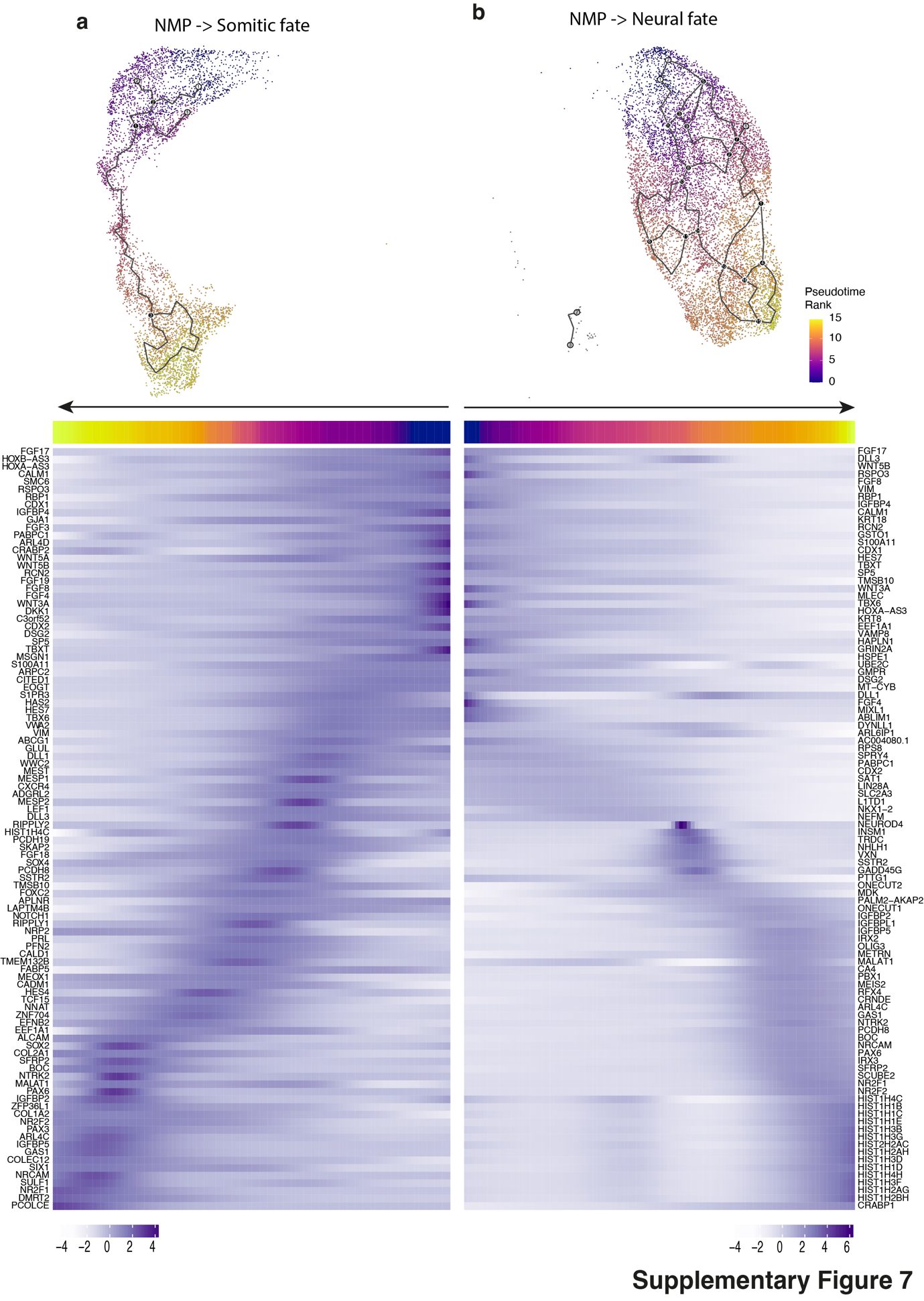

**Supplementary Figure 7: Neural and mesodermal trajectory gene expression signatures.** Monocle trajectory analysis showing two lineages from neuromesodermal progenitor (NMP) population towards somitic (a) and neural (b) populations (top), alongside ordering of cells by pseudotime with heatmap representation of the top 100 variable genes for each differentiation route (bottom).

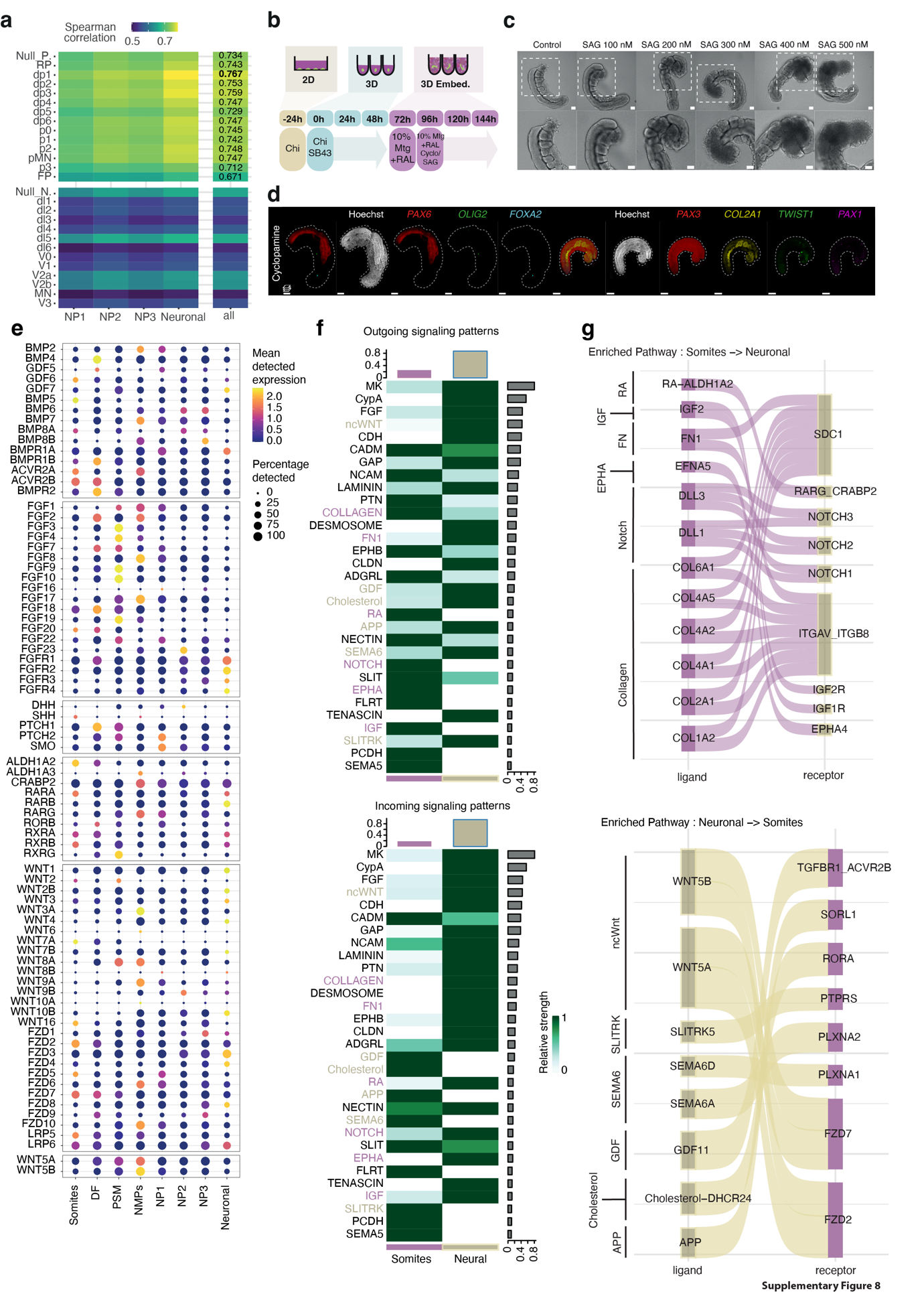

**Supplementary Figure 8: Dorsoventral patterning and CellChat analysis in hTLS.** (**a**) Correlation coefficients between neural populations in 120h hTLS and the human embryonic neural tube^68^. NP, Neural progenitor; Null_P, unannotated progenitor; RP, Roof plate; dp, Dorsal progenitor; p, Progenitor; pMN, Motor Neuron progenitor; FP, Floor plate. (**b)** Schematic diagram of hTLS treatment with SAG (sonic hedgehog (SHH) pathway activator) and cyclopamine (SHH pathway inhibitor). Chi, Chrion (CHIR99021); Mat, Matrigel; SB43, SB431542; RAL, retinal. (**c)** Representative brightfield images of SAG-treated hTLSs at 144h. Dashed box represents position of enlarged region. Scale bars, 100 μm. (**d)** Projected images of HCR staining in 144h hTLSs treated with cyclopamine. Dashed outline hTLSs. Scale bars, 100 µm. (**e**) Gene expression of signalling ligands and receptors at single cell level across populations. (**f**) CellChat analysis showing statistically significant interactions between the somite and neuronal populations. Pathway names coloured in purple indicate possible somite-to-neural directional signalling, and names coloured in ochre indicate possible neural-to-somite directional signalling. (**g**) CellChat-predicted interactions between ligands and receptors, organised by pathway.

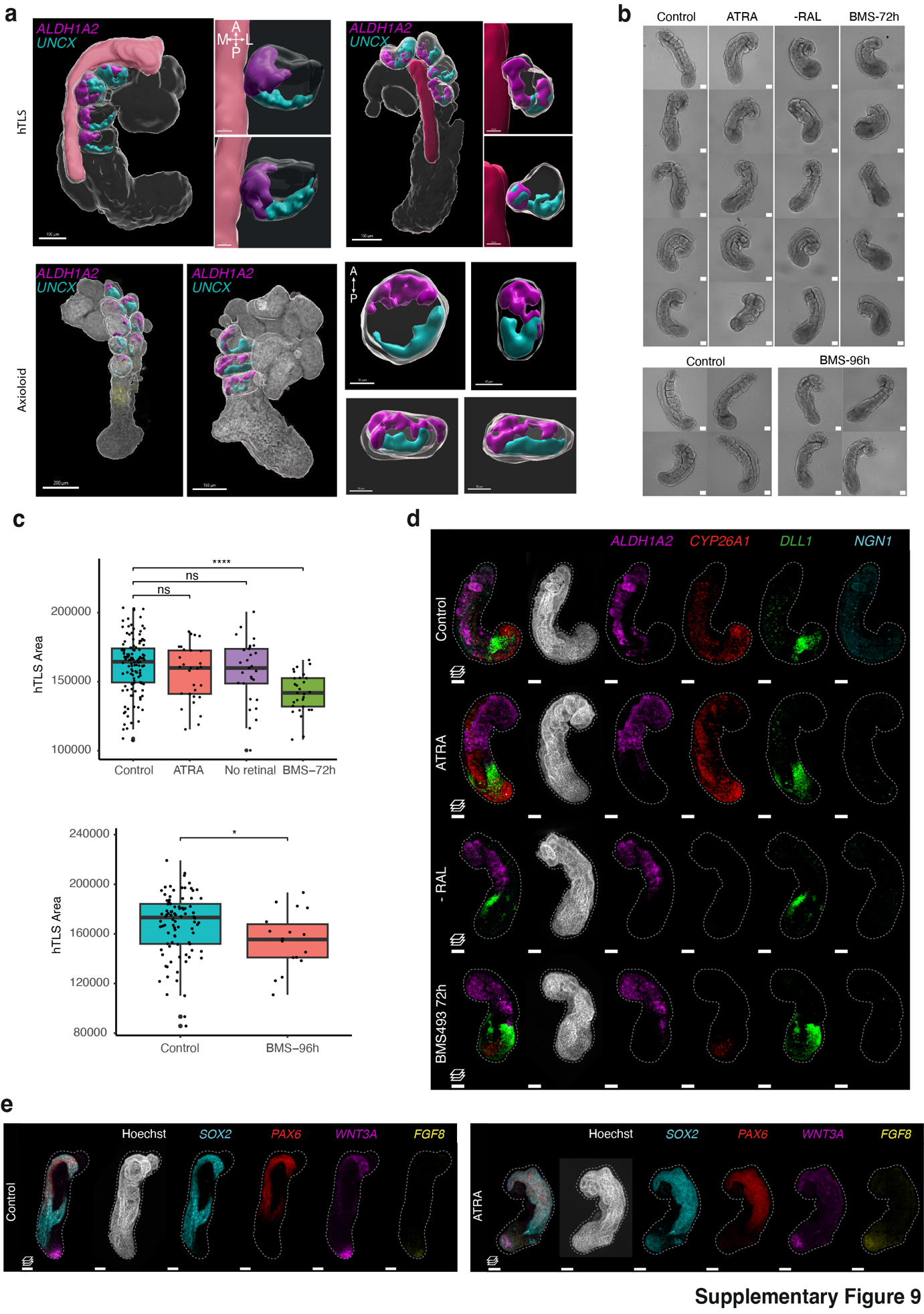

**Supplementary Figure 9: Signalling interactions between somites and neural tube in hTLS models. (a)** Imaris-rendered surfaces showing regions of *ALDH1A2* (purple) and *UNCX* (blue) expression by HCR within somites, relative to the position of the neural tube (pink), for both hTLS and axioloid structures. A, anterior; P, posterior; M, medial; L, lateral. (**b**) Representative brightfield images of 120h hTLS with replacement of RAL with all *trans*-retinoic acid (ATRA), without retinal (RAL), or with treatment with BMS493 (RA inhibitor) at 72h (top) and 96h (bottom). Scale bars, 100 µm. * indicates *P* < 0.05 and **** indicates *P* < 0.0001 obtained by Wilcoxon test. **(c)** Area quantification of hTLS on RA signalling pathway modulation. **(d)** Projected images of 120h hTLS HCR-stained following RA signalling pathway modulation. Scale bars, 100 μm. Dashed lines outline hTLS. (**e)** Projected images of 120h hTLS HCR-stained following RA signalling pathway modulation. Scale bars, 100 μm. Dashed lines outline hTLS. RA, Retinoic acid; ATRA, all-trans Retinoic Acid; RAL, Retinal.

**Supplementary Video 1: ZO-1 expression in hTLS.** Live-cell imaging of ZO-1 human iPSC cell line, showing widefield and ZO-1::mEGFP expression in hTLS from 72.5-96 hours. Scale bars, 100 μm.

**Supplementary Video 2: HES7 oscillatory expression in hTLS.** Live-cell imaging of HES7::Achillies human iPSC cell line, showing HES7 expression in hTLS from 100-121 hours**.** Scale bars, 100 μm.

**Supplementary Video 3: *ALDH1A2* expression in somites of mouse embryos.** Localised *ALDH1A2* expression found mediolateral in somites to the neural tube. Scale bars, 200 μm.
